# Ribosome profiling reveals the fine-tuned response of *Escherichia coli* to mild and severe acid stress

**DOI:** 10.1101/2023.06.02.543275

**Authors:** Kilian Schumacher, Rick Gelhausen, Willow Kion-Crosby, Lars Barquist, Rolf Backofen, Kirsten Jung

## Abstract

The ability to respond to acidic environments is crucial for neutralophilic bacteria. *Escherichia coli* has a well-characterized regulatory network that triggers a multitude of defense mechanisms to counteract excess of protons. Nevertheless, systemic studies of the transcriptional and translational reprogramming of *E. coli* to different degrees of acid stress have not yet been performed. Here, we used ribosome profiling and RNA sequencing to compare the response of *E. coli* (pH 7.6) to sudden mild (pH 5.8) and severe near-lethal acid stress (pH 4.4) conditions that mimic passage through the gastrointestinal tract. We uncovered new differentially regulated genes and pathways, key transcriptional regulators, and 18 novel acid-induced candidate sORFs. By using machine learning leveraging large compendia of publicly available *E. coli* expression data, we were able to distinguish between the response to acid stress and general stress. These results expand the acid resistance network and provide new insights into the fine-tuned response of *E. coli* to mild and severe acid stress.

## Introduction

The infective dose of enteropathogens varies significantly among bacterial genera, and is dependent on the number and complexity of acid resistance mechanisms (see Schwarz *et al.*, 2022 for review).^1^ *Escherichia coli* is equipped with a high number of defense mechanisms to survive the acidity of the stomach and correspondingly can have an infective dose of as low as less than 50 cells.^2^ Enterobacteria that survive the stomach also confront mild acid stress in the colon, due to the presence of short-chain fatty acids produced by obligate anaerobes.^3^ Other neutralophilic bacteria encounter low pH environments in a variety of settings, such as acidic soils, fermented food, or phagosomes within macrophages.^1^

The cytoplasmic membrane represents a primary barrier for protons (H^+^). Nevertheless, at low pH, H^+^ can permeate into the cytoplasm via protonated water chains, ion channels, or damaged membranes.^4^ Upon acidification, the cytoplasmic pH transiently decreases but returns to neutral within short time intervals. For example, a reduction in external pH to 5.5 causes a temporary decline in internal pH to ∼6.0.^5^ Intracellular acidification leads to protonation of biological molecules, ultimately altering their charge and structure. This, in turn, may lead to protein unfolding, denaturation and reduced enzymatic activities.^6^ Furthermore, acidification causes membrane and DNA damage.^7, 8^

To maintain a constant internal pH and balance fluctuations in H^+^ concentrations, the intrinsic buffering capacity of the cytoplasm is crucial, as protons can be sequestered by side-chains of proteins, inorganic phosphates, polyphosphates, or polyamines.^9^ Additional protective mechanisms that counteract acid stress and ensure survival in low pH habitats involve proton pumps, membrane remodeling, acid-dependent chemotaxis, chaperones, acid shock proteins and the induction of enzyme-based H^+^-consuming acid resistance (AR) systems.^10, 11^

*E. coli* is equipped with four different AR systems, namely the glutamate decarboxylase (Gad, AR2), arginine decarboxylase (Adi, AR3), lysine decarboxylase (Cad, AR4) and ornithine decarboxylase (Orn, AR5) systems.^12, 13^ Each AR system consists of an H^+^-consuming amino acid decarboxylase (GadA/GadB, AdiA, CadA, SpeF) and a corresponding antiporter (GadC, AdiC, CadB, PotE), which serves to uptake the amino acid and export the more alkaline reaction product into the surrounding medium. This strategy ensures a simultaneous increase in intracellular and extracellular pH.^7, 13, 14^ Notably, each AR system is activated at different external pH values and growth phases. We have previously found that individual *E. coli* cells exposed to consecutively increasing acid stress activate the Gad, Adi and Cad system heterogeneously, resulting in functional diversification and enhanced population fitness.^15, 16^ The regulatory network of the Gad system is highly complex and involves more than ten regulatory elements,^17^ whereas the Adi and Cad systems are each regulated by a single transcriptional regulator, AdiY and CadC, respectively.^16^ Nevertheless, additional regulatory elements seem to exist that link not only these three ARs in individual cells, but also contribute to a fine-tuned response of *E. coli* to different levels of acid stress^16^.

Previous systematic studies to determine genome-wide adaptations within the acid stress response of *E. coli* have either been restricted to microarrays,^18^ two-dimensional gel electrophoresis,^19, 20^ or have only been performed under mild acidic conditions (pH 5.0 – 6.0).^21–, 23^ Here, we present the first systemic study of global adaptations to different intensities of acid stress in *E. coli* at both the transcriptional and translational levels. Specifically, we compare gene expression and translation at pH 5.8 and pH 4.4, with pH 7.6, using RNA sequencing (RNA-Seq) and ribosome profiling (Ribo-Seq). Ribo-Seq allows deep-sequencing of ribosome protected mRNA fragments (RPFs),^24, 25^ which are obtained by nuclease digestion of non-ribosome covered RNA, and has significantly advanced the current understanding of translational regulation.^26^ The applications of Ribo-Seq are manifold, and include monitoring protein synthesis rates across the proteome, identification of novel small open reading frames (sORFs), as well as determining protein copy numbers per cell during steady-state growth.^27, 28^ In contrast to mass spectrometry-based approaches, ribosome profiling is independent of protein biochemistry and allows detection of small proteins with less than 50 amino acids in length.^29^ In recent years, substantial advances in understanding translational events have been made through the utilization of Ribo-Seq, not only in various bacterial species, but also in archaea, bacteriophages and microbiomes.^28, 30–35^

Our results reveal hundreds of differentially transcribed and translated genes of *E. coli* K-12 at pH 5.8 and 4.4, as well as examples of pH-dependent changes in translation efficiency. We identified previously undiscovered biological processes in response to acidity including increased siderophore synthesis, glycerol catabolism, copper efflux, nucleotide biosynthesis, and spermidine/multidrug export as well as decreased membrane transport and metabolic activities. In addition, we have identified new transcription factors (TFs) as key players during low pH exposure, and 18 novel candidate sORFs involved in the response to acid stress. Finally, we differentiated acid-specific transcriptional adaptations by using machine learning to compare the low pH response to that of other stressors.

## Results and Discussion

### 1. Examination of alterations in translatome and transcriptome of *E. coli* in response to varying degrees of acid stress

To mimic natural stress conditions, such as passage of *E. coli* through the gastrointestinal tract, we established the following protocol involving a sudden change to low pH, detection of a rapid response, and severe, near-lethal acid stress. Specifically, *E. coli* K-12 MG1655 was cultivated in unbuffered lysogeny broth (LB medium) at pH 7.6 until exponential growth phase (OD_600_ = 0.5). Then, 5 M hydrochloric acid was added directly to expose the cells to a pH of 5.8 or stepwise to a pH of 4.4, corresponding to mild and severe acid stress (Fig. 1A). The final optical densities were comparable (pH 7.6: OD_600_ = ∼1.1; pH 4.4: OD_600_ = ∼ 0.7) (Table S1) and the pH values hardly changed compared to t_0_ (Fig. 1A, Table S2). To investigate whether a 15-minute exposure to pH 4.4 was sufficient to induce cellular adaption to severe acid stress, we examined the temporal dynamics of *adiA* expression by RT-qPCR. We detected an increase in *adiA* mRNA levels as early as 15 minutes after the shift to pH 4.4, and no substantial further increase after 30 or 60 minutes (Fig. S1). Additionally, we evaluated cell viability at the final experimental time points using propidium iodide staining.^36^ The percentage of dead cells detected was less than 1% at pH 7.6, 5% at pH 5.8, and 18% at pH 4.4 (Fig. S2).

**Figure 1:**
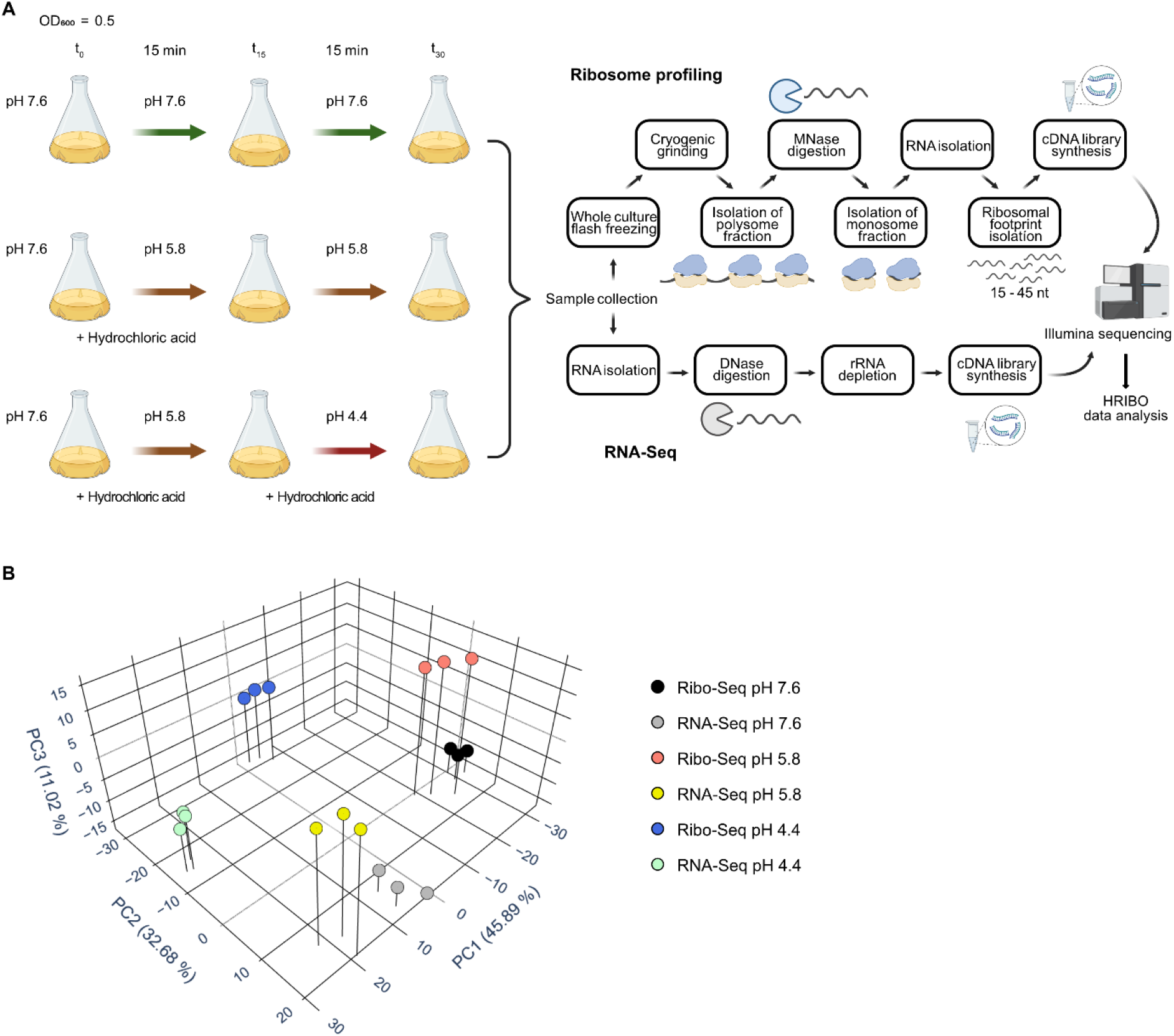
Schematic overview of culture conditions, pH-shift procedures, and the Ribo-Seq and RNA-Seq workflow. **(A)** Biological triplicates of *E. coli* MG1655 cells were grown in unbuffered LB medium (pH 7.6) to an OD_600_ = 0.5 (t_0_). Subsequently, the cultures were either grown for an additional 30 minutes at pH 7.6, shifted for 30 minutes to pH 5.8, or shifted first for 15 minutes to pH 5.8 (t_15_) and then to pH 4.4 (t_30_). pH-shifts were initiated by direct addition of 5 M HCl to the cultures. At t_30_ samples were collected. For RNA-Seq, total RNA was isolated, DNase digested, and ribosomal RNA was depleted prior to cDNA library preparation. For Ribo-Seq, whole cultures were flash-frozen in liquid nitrogen and cryogenically grinded using a Freezer Mill. Polysome fractions were isolated, and non-ribosome protected RNA was digested using MNase. Ribosomal footprints were purified from the monosome fraction using a size selection gel and converted to cDNA libraries. Upon Illumina sequencing, data were analyzed using the HRIBO pipeline.^40^ **(B)** Principal component analysis (PCA) of median-of-ratios normalized and regularized log-transformed (rlog) read count values for *E. coli* Ribo-Seq and RNA-Seq triplicate data at pH 4.4, 5.8 and 7.6. Panel (A) was created with BioRender.com.

Next, cells were harvested and lysed as previously described by whole culture flash freezing and cryogenic grinding in a freezer mill to avoid bias from translation-arresting drugs and filtering.^37^ The subsequent steps of our ribosome profiling protocol were a combination of methodologies reported by Latif *et al.*^38^ and Mohammad & Buskirk^39^ (see Methods section for details). Strand-specific Illumina sequencing yielded an average of approximately 30 million cDNA reads per sample for Ribo-Seq and 5-10 million for RNA-Seq. The next generation sequencing data was analyzed using an extended version of the high-throughput HRIBO data analysis pipeline.^40^ All samples achieved sufficient coverage with over two million reads each mapping uniquely to the coding regions. The rRNA contamination was higher in the pH 4.4 Ribo-Seq samples compared with other conditions but accounted for less than 15 % in all cDNA libraries (Fig. S3). The length distribution of the generated ribosome protected fragments (RPFs) was broad, ranging from 15 to 45 nucleotides (Fig. S4), consistent with previous observations in other prokaryotic ribosome profiling analyses.^37, 38^ We did not detect stress-induced ribosome accumulation in the initiation region of ORFs, contrary to previous observations in *E. coli* during heat stress^41^ and in yeast during oxidative stress.^42^ In fact, ribosome occupancy in the translation initiation regions was slightly reduced at pH 4.4 and 5.8 compared with physiological pH (Fig. S5). This could be explained due to diminished ribosome-RNA complex stability and increased ribosome drop-off under acidic conditions. The biological triplicates for each experimental condition clustered on the first three principal components in a PCA plot (Fig. 1B). Notably, the global gene expression profiles were highly distinct at pH 4.4 compared with both pH 5.8 and 7.6.

### 2. Coordinated regulation of transcription and translation in response to acid stress

The tool *deltaTE*^43^ was used to assess transcriptional and translational changes (i.e., differential expression and differential translation efficiency) in response to mild and severe acid stress. Low-expression transcripts were filtered out, and we focused our analysis on 3,654 genes with mean reads per kilobase per million reads mapped (rpkm) values ≥ 5 across all investigated conditions. Our findings reveal that 702 transcripts were significantly altered at pH 5.8 compared with physiological pH (absolute mRNA log_2_FC ≥ 1 and FDR (False discovery rate) adjusted p-values ≤ 0.05), and 1,030 genes showed significant differences in mRNA levels at pH 4.4 (Fig. 2A). These results suggest that extensive transcriptional reprogramming occurred, which was influenced by the degree of acid stress. As illustrated by the Venn diagram overlaps (Fig. 2), a large number of adaptations occurred regardless of the degree of acid stress. Nonetheless, several hundred genes were differentially expressed exclusively at pH 5.8 or 4.4 (Fig. 2A). This suggests that in addition to universal adaptations at low pH, specific adaptions for mild and severe acid stress occur. We further determined the number of genes with stress-dependent alterations in ribosome-protected-fragment (RPF) counts to be 679 at pH 5.8 and 1,440 at pH 4.4 (absolute RPF log_2_FC ≥ 1 and FDR adjusted p-values ≤ 0.05), which was in a similar range compared with the RNA-Seq data (Figs. 2A, 2B). Accordingly, the global FC values for mRNA and RPF levels showed a high Pearson correlation coefficient (r) under both conditions (Figs. 2C, 2D, gray dots). This indicates that transcriptional regulation of these genes is the predominant response to acid stress. However, a subset of genes exhibited exclusive significant regulation at either the transcriptional (red dots) or translational (blue dots) level.

**Figure 2:**
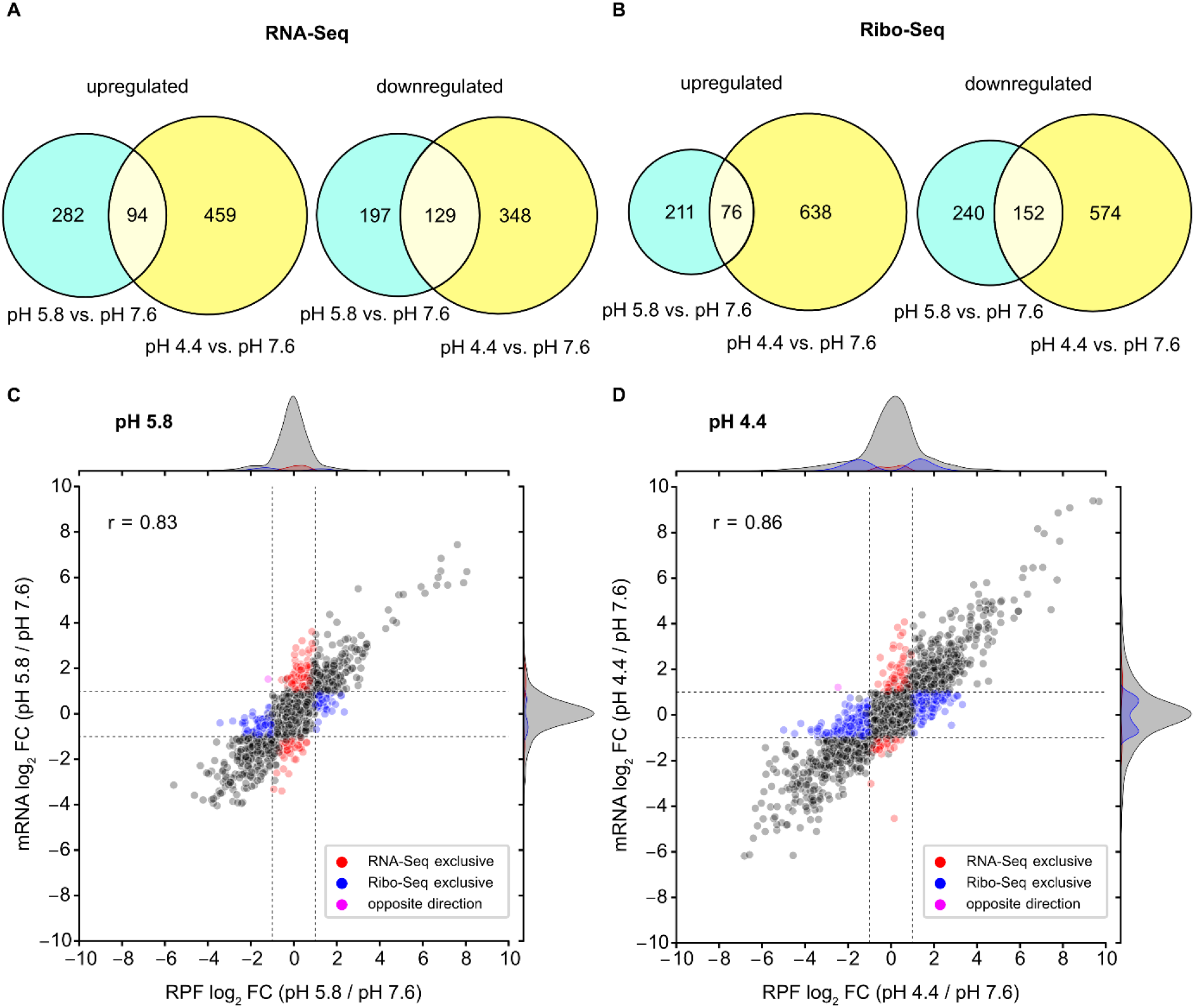
Genome-wide adaptations correlate at the transcriptional and translational levels in *E. coli* under acid stress. Weighted Venn diagrams show the total number and overlap of genes with significant FCs (absolute log_2_ FC ≥ 1 and p-adjust ≤ 0.05) determined by **(A)** RNA-Seq or **(B)** Ribo-Seq for cells exposed to pH 5.8 or pH 4.4, compared to the control (pH 7.6). **(C)** Comparison of global RPF and mRNA log_2_ FC values for pH 5.8, or **(D)** pH 4.4 vs. pH 7.6. Dashed lines indicate log_2_ fold change values of +1 or −1. Hundreds of genes exhibited differential expression (absolute log_2_ FC ≥ 1 and p-adjust ≤ 0.05) at both the transcriptional and translational levels, whereas others were exclusively detected by either RNA-Seq (red dots) or Ribo-Seq (blue dots), or had significant changes in opposite directions (pink dots). Values of the Pearson correlation coefficient (r) are indicated. FC, fold change; RPF, ribosome-protected-fragments.

Specifically, at pH 5.8, 193 genes were detected to be significantly regulated exclusively by RNA-Seq, while 216 genes were exclusively affected in the Ribo-Seq data (Fig. 2C, Table S3). At pH 4.4, 127 differentially regulated genes were found exclusively by RNA-Seq and 570 genes by Ribo-Seq (Fig. 2D, Table S3). Notably, for *fruA* at pH 5.8 and *yecH* at pH 4.4, opposite changes were observed at the transcriptional and translational levels (Figs. 2C, 2D, pink dots).

Next, we investigated translation efficiency (TE) to identify genes that undergo translational regulation in response to acidic conditions. TE provides information regarding ribosome counts per mRNA and is calculated as the ratio of RPFs over transcript counts within a gene’s coding sequence normalized to mRNA abundance.^43^ We identified 22 genes at pH 5.8 and 89 genes at pH 4.4, which displayed significantly altered TEs (absolute log_2_ TE fold change ≥ 1 and p-adjust ≤ 0.05) (Table S4). The highest increase in TE at pH 4.4 was found for the KpLE2 phage-like element (*topAI*), a hydroxyethylthiazole kinase (*thiM*), and a palmitoleoyl acyltransferase (*lpxP*). In contrast, *yecH* and *yjbE*, both encoding uncharacterized proteins, and *malM* of the maltose regulon, showed the most prominent decrease in TE at pH 4.4 (Table S4). At pH 5.8, we noted the largest increase in TE for a ferredoxin-type protein encoded by *napF*, an iron transport protein (*feoA*) and a tagaturonate reductase (*uxaB*). Conversely, the largest decrease was observed for a protein of the fructose-specific phosphotransferase system (*fruA*), a tripartite efflux pump membrane fusion protein (*emrK*) and an HTH-type transcriptional regulator (*ydeO*) (Table S4).

In summary, besides extensive transcriptional reprogramming, dozens of genes exhibit significant FCs either at the transcriptional or translational level in response to acid stress. This underlines that transcription and translation are not always coupled in bacteria. Similar findings were reported by Zhang et al. 2017, who conducted Ribo-Seq and RNA-Seq analysis for *E. coli* under heat stress.^41^ Overall, such differential regulation can be explained, for example, by delayed translation relative to transcript synthesis, selective recruitment or release of ribosomes, or regulation during translation initiation, elongation, or ribosome biogenesis,^44–48^ which could be beneficial under stress conditions.

### 3. Functional implications of genes with differential mRNA and RPF levels under mild acid stress

To obtain a more profound understanding of the fine-tuned response of *E. coli* to different degrees of acid stress, we first analyzed all genes with differential mRNA and ribosome coverage levels during mild acid stress (pH 5.8). Under this condition, the top candidates with the highest FC values for mRNA and RPF are: (i) the *cad* operon, encoding the core components of the Cad AR system (see Chapter 5), (ii) the *glp* regulon, responsible for glycerol and *sn*-glycerol 3-phosphate uptake and catabolism,^49^ (iii) the *mdtJI* operon, encoding a heterodimeric multidrug/spermidine exporter,^50^ and (iv) genes encoding proteins involved in motility and flagella biosynthesis (Table 1). A comprehensive list of normalized read counts, mRNA and RPF FCs, and TEs for all *E. coli* genes is provided in Table S5. We tested a representative selection of differentially expressed genes by RT-qPCR. In all cases, the detected changes in mRNA levels were consistent with the data gathered by RNA-Seq (Fig. S6).

**Table 1:**
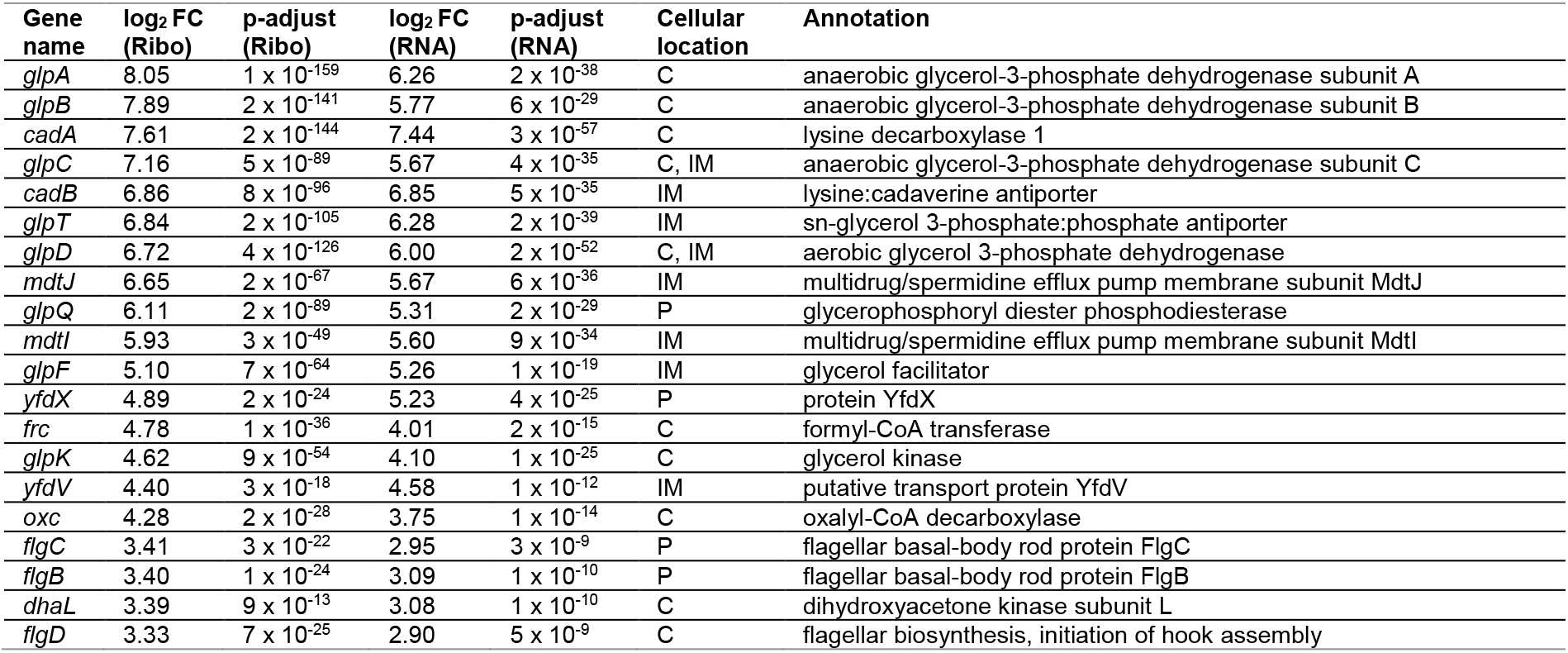
Top 20 genes with increased RPF levels at pH 5.8 compared to pH 7.6, sorted in descending order by Ribo-Seq log_2_ FC values. C, cytosol; P, periplasm; IM, inner membrane.

| Gene name | log<sub>2</sub> FC (Ribo) | p-adjust (Ribo) | log<sub>2</sub> FC (RNA) | p-adjust (RNA) | Cellular location | Annotation |
| --- | --- | --- | --- | --- | --- | --- |
| *glpA* | 8.05 | 1 x 10<sup>-159</sup> | 6.26 | 2 x 10<sup>-38</sup> | C | anaerobic glycerol-3-phosphate dehydrogenase subunit A |
| *glpB* | 7.89 | 2 x 10<sup>-141</sup> | 5.77 | 6 x 10<sup>-29</sup> | C | anaerobic glycerol-3-phosphate dehydrogenase subunit B |
| *cadA* | 7.61 | 2 x 10<sup>-144</sup> | 7.44 | 3 x 10<sup>-57</sup> | C | lysine decarboxylase 1 |
| *glpC* | 7.16 | 5 x 10<sup>-89</sup> | 5.67 | 4 x 10<sup>-35</sup> | C, IM | anaerobic glycerol-3-phosphate dehydrogenase subunit C |
| *cadB* | 6.86 | 8 x 10<sup>-96</sup> | 6.85 | 5 x 10<sup>-35</sup> | IM | lysine:cadaverine antiporter |
| *glpT* | 6.84 | 2 x 10<sup>-105</sup> | 6.28 | 2 x 10<sup>-39</sup> | IM | sn-glycerol 3-phosphate:phosphate antiporter |
| *glpD* | 6.72 | 4 x 10<sup>-126</sup> | 6.00 | 2 x 10<sup>-52</sup> | C, IM | aerobic glycerol 3-phosphate dehydrogenase |
| *mdtJ* | 6.65 | 2 x 10<sup>-67</sup> | 5.67 | 6 x 10<sup>-36</sup> | IM | multidrug/spermidine efflux pump membrane subunit MdtJ |
| *glpQ* | 6.11 | 2 x 10<sup>-89</sup> | 5.31 | 2 x 10<sup>-29</sup> | P | glycerophosphoryl diester phosphodiesterase |
| *mdtI* | 5.93 | 3 x 10<sup>-49</sup> | 5.60 | 9 x 10<sup>-34</sup> | IM | multidrug/spermidine efflux pump membrane subunit MdtI |
| *glpF* | 5.10 | 7 x 10<sup>-64</sup> | 5.26 | 1 x 10<sup>-19</sup> | IM | glycerol facilitator |
| *yfdX* | 4.89 | 2 x 10<sup>-24</sup> | 5.23 | 4 x 10<sup>-25</sup> | P | protein YfdX |
| *frc* | 4.78 | 1 x 10<sup>-36</sup> | 4.01 | 2 x 10<sup>-15</sup> | C | formyl-CoA transferase |
| *glpK* | 4.62 | 9 x 10<sup>-54</sup> | 4.10 | 1 x 10<sup>-25</sup> | C | glycerol kinase |
| *yfdV* | 4.40 | 3 x 10<sup>-18</sup> | 4.58 | 1 x 10<sup>-12</sup> | IM | putative transport protein YfdV |
| *oxc* | 4.28 | 2 x 10<sup>-28</sup> | 3.75 | 1 x 10<sup>-14</sup> | C | oxalyl-CoA decarboxylase |
| *flgC* | 3.41 | 3 x 10<sup>-22</sup> | 2.95 | 3 x 10<sup>-9</sup> | P | flagellar basal-body rod protein FlgC |
| *flgB* | 3.40 | 1 x 10<sup>-24</sup> | 3.09 | 1 x 10<sup>-10</sup> | P | flagellar basal-body rod protein FlgB |
| *dhaL* | 3.39 | 9 x 10<sup>-13</sup> | 3.08 | 1 x 10<sup>-10</sup> | C | dihydroxyacetone kinase subunit L |
| *flgD* | 3.33 | 7 x 10<sup>-25</sup> | 2.90 | 5 x 10<sup>-9</sup> | C | flagellar biosynthesis, initiation of hook assembly |

Next, we performed gene set enrichment analysis (GSEA) using *clusterProfiler*^51^ to identify biological processes associated with differentially expressed genes (DEGs) at pH 5.8. Among the most enriched Gene Ontology (GO) terms for biological processes at pH 5.8 was ‘spermidine transmembrane transport’ (Fig. 3), which corresponds to the induction of *mdtJI* (Table 1) and a polyamine ABC transporter encoded by *potABCD* (Table S5). Polyamines are crucial for survival under acid stress, as they reduce membrane permeability by blocking OmpF and OmpC porins.^52–54^ External spermidine supplementation also improved acid resistance in *Streptococcus pyogenes*.^55^ On the other hand, overaccumulation of polyamines can be toxic and potentially lethal for *E. coli*.^50, 56^ Therefore, precise transmembrane transport of polyamines in acidic environments is critical and contributes to survival in acidic conditions.

**Figure 3:**
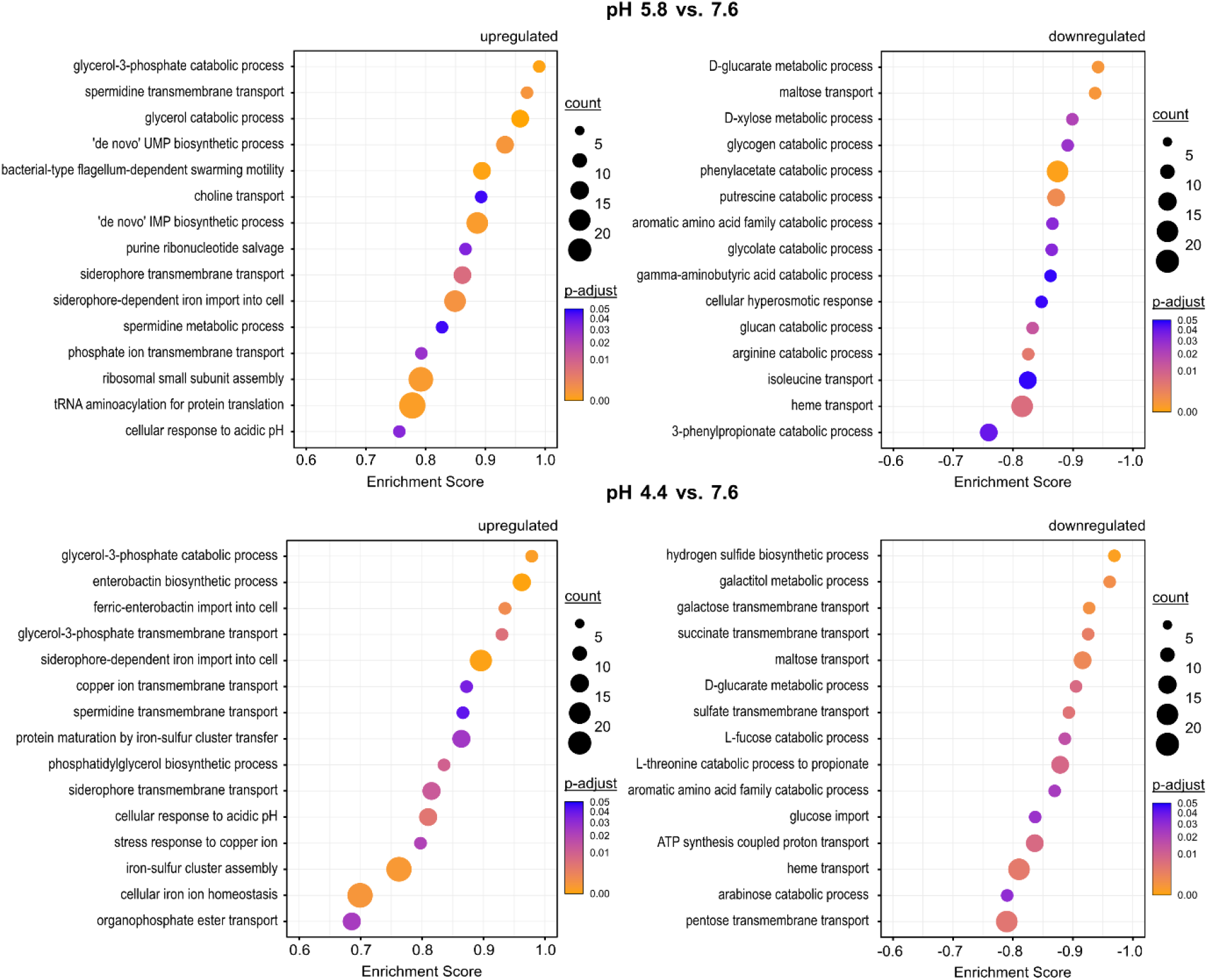
Up- and downregulated biological processes in *E. coli* at pH 5.8 and 4.4. Gene Set Enrichment Analysis (GSEA) was conducted using the *gseGO* function in the *clusterProfiler* package^51^ with the ribosome profiling differential expression data sorted by log_2_ fold change values as input. GO terms were considered as either up- or downregulated if p-adjust values were ≤ 0.05. The top 15 non-redundant GO terms were sorted in descending order by the *clusterProfiler* Enrichment Score, and are shown for pH 5.8 vs. 7.6, and pH 4.4 vs. 7.6. The dot size represents the number of genes associated with each GO term, the dot color represents adjusted p-values corrected for the false discovery rate.

The enrichment of the GO terms ‘glycerol-3-phosphate catabolic process’ and ‘glycerol catabolic process’ at pH 5.8 (Fig. 3) has not yet been associated with acid stress to our knowledge. Notably, of the 14 genes with the largest increase in RPF counts at pH 5.8, seven belong to the *glp* regulon (Table 1). This regulon is required for the uptake and catabolism of glycerol and *sn*-glycerol 3-phosphate (G3P).^49^ In this pathway, G3P is converted to dihydroxyacetone phosphate by membrane-bound dehydrogenases, either aerobically via GlpD or anaerobically by the GlpABC complex.^57, 58^ Alternatively, dihydroxyacetone phosphate can be produced directly from glycerol by GldA and the protein products of the *dhaKLM* operon.^59^ The *dhaKLM* operon was also induced at pH 5.8 (Table S5). It remains unclear whether glycerol and G3P catabolism directly contribute to acid tolerance or whether the *glp* regulon is activated as a consequence of other low pH adjustments. Expression of *glp* genes is regulated by the repressor GlpR, which is inactivated upon binding of glycerol or G3P.^60^ We hypothesize that changes in phospholipid composition under acid stress conditions^61^ may release G3P, which in turn induces the *glp* regulon. Accordingly, the GO term ‘phosphatidylglycerol biosynthetic process’ was enriched under acid stress (Fig. 3).

Another observation is the upregulation of *de novo* biosynthesis pathways for pyrimidine and purine nucleotides at pH 5.8 (Fig. 3). The induction of a large proportion of the PurR-dependent regulon involved in *de novo* nucleotide synthesis (Fig. 3, Table S5) suggests that *E. coli* requires additional nucleotides to cope with the extensive transcriptional reprogramming. Besides, intracellular acidification can lead to DNA damage, such as depurination,^62^ making enhanced nucleotide biosynthesis a critical compensatory mechanism. Recently, *Oenococcus oeni* was reported to experience a decrease in the abundance of both purines and pyrimidines under acid stress, while nucleotide metabolism and transport increased,^63^ suggesting a similar phenomenon in this species. Other enriched GO terms under mild acid stress include ‘choline transport’, ‘siderophore transmembrane transport’, ‘phosphate ion transmembrane transport’, ‘ribosomal small subunit assembly’, ‘tRNA aminoacylation for protein translation’ and ‘bacterial-type flagellum-dependent swarming motility’ (Fig. 3). pH-dependent motility has previously been observed in *E. coli*, *Salmonella*, and *Helicobacter* (reviewed in Schwarz et al., 2022).^1^ These observations suggest that bacterial cells use an escape strategy to migrate to more favorable pH environments when challenged with acidic conditions.

On the contrary, our findings reveal that many membrane and periplasmic proteins (18 of the 20 genes with the most diminished RPF counts, Table 2) were among the top candidates with decreased mRNA and RPF levels under mild acid stress. This affected, for example, genes encoding ABC transporters (*mal* regulon, *dpp* operon) and symporters (*actP*, *melB*, *gabP*), highlighting the superiority of Ribo-Seq over mass spectrometry-based approaches, namely its independence of protein biochemistry and higher sensitivity.^29^ Furthermore, GSEA identified membrane transport and metabolic activities as the most downregulated biological processes in response to mild acid stress. For example, ‘maltose transport’, ‘isoleucine transport’, ‘heme transport’, ‘putrescine catabolic process‘, ‘glycolate catabolic process’ and ‘aromatic amino acid family catabolic process’ were among the most downregulated GO terms at pH 5.8 (Fig. 3). Downregulation of H^+^-coupled transport processes represents a key mechanism by which *E. coli* restricts proton influx into cells. In addition, the downregulated metabolic processes are in many cases associated with the synthesis and conversion of amino acids and carbon sources. For example, the catabolism of aromatic amino acids and arginine was also reduced at pH 5.8 (Fig. 3). Particularly noteworthy is the downregulation of the arginine catabolic pathway, which involves the protein products of the *astEBDAC* operon. At pH 5.8, hardly any reads mapped in the *astEBDAC* region, despite detectable expression at pH 7.6 and pH 4.4 (Table S5). Presumably, *E. coli* preserves the intracellular arginine pool at pH 5.8, as this amino acid serves as a substrate for the Adi system during severe acid stress.^16, 64^

**Table 2:**
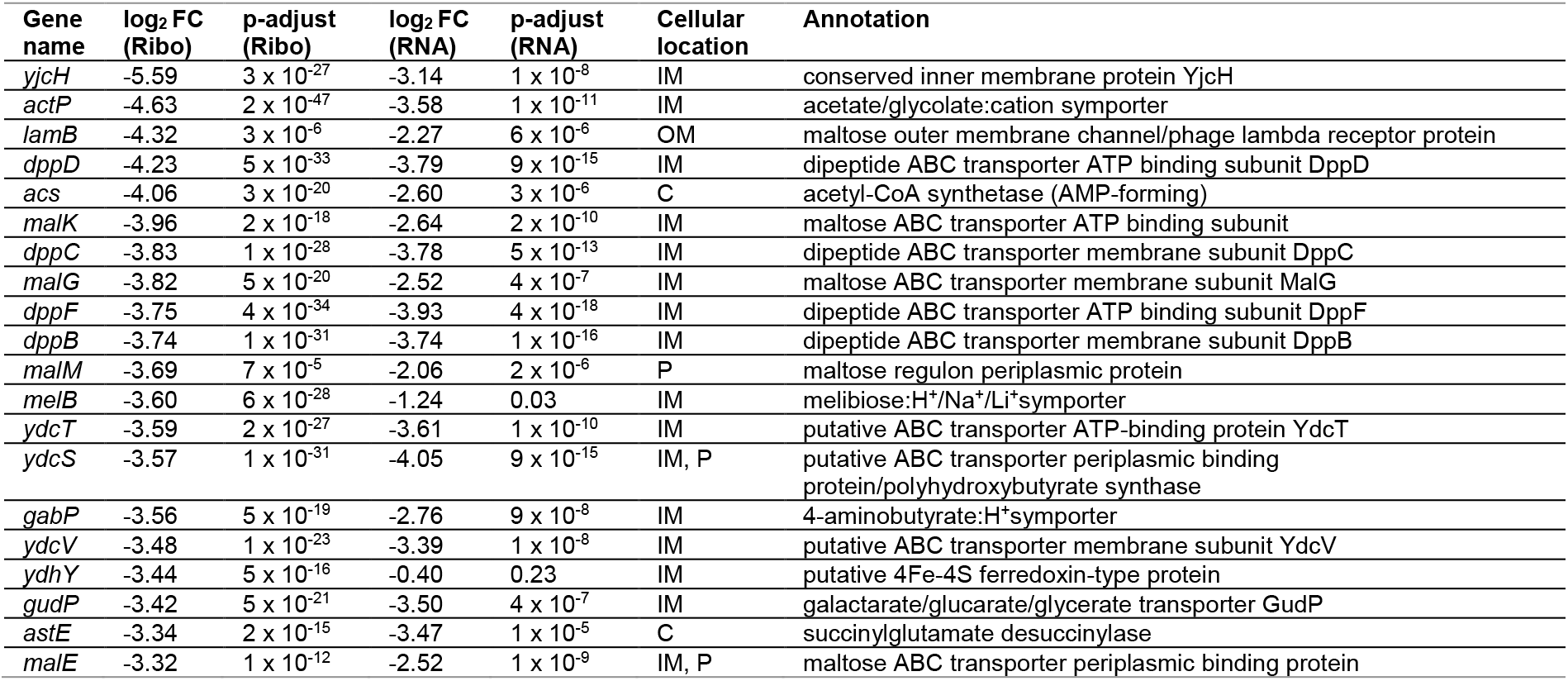
Top 20 genes with decreased RPF levels at pH 5.8 compared to pH 7.6, sorted in ascending order by Ribo-Seq log_2_ FC values. C, cytosol; P, periplasm; IM, inner membrane; OM, outer membrane.

In summary, the response of *E. coli* to mild acid stress is characterized by the activation of the motility machinery to escape to less acidic habitats, by induction of the *cad* operon, and genes involved in polyamine transport and glycerol-3-phosphate conversion (Tables 1 and 2, Fig. 3). In addition, *E. coli* restricts the influx of protons and conserves energy by reducing its metabolic activities.

### 4. Functional implications of genes with differential mRNA and RPF levels under severe acid stress

Next, we analyzed genes with differential mRNA and ribosome coverage levels in response to severe acid stress (pH 4.4) compared with nonstress (pH 7.6). Genes with the highest number of increased read counts, which were not already upregulated at pH 5.8, were *asr,* encoding an acid shock protein, followed by *bdm,* encoding a biofilm-modulation protein, and *bhsA,* encoding a multiple stress resistance outer membrane protein (Table 3). Originally, Asr was classified as a periplasmic acid shock protein, although its role in acid adaptation remained unclear.^65^ Recently, Asr was shown to be an intrinsically disordered chaperone that contributes to outer membrane integrity and to act as an aggregase in order to prevent aggregation of proteins with positive charges.^66^ Our Ribo-seq data clearly illustrate the enormous importance of Asr under severe acid stress in *E. coli*, as it is one of the most abundant proteins in the cell, with approximately 2% of all reads mapping in the *asr* coding region at pH 4.4 (corresponding to a ∼1000-fold upregulation compared to pH 7.6). Strikingly, almost half of the top 20 genes with increased ribosome coverage of transcripts (*ydgU*, *yhcN*, *yjcB*, *yedR*, *yhdV*, *ybiJ*, *ycgZ* and *ycfJ*) are poorly characterized (Table 3). So far, only YhcN from the above list has been shown to be involved in the response to acid stress.^67^

**Table 3:**
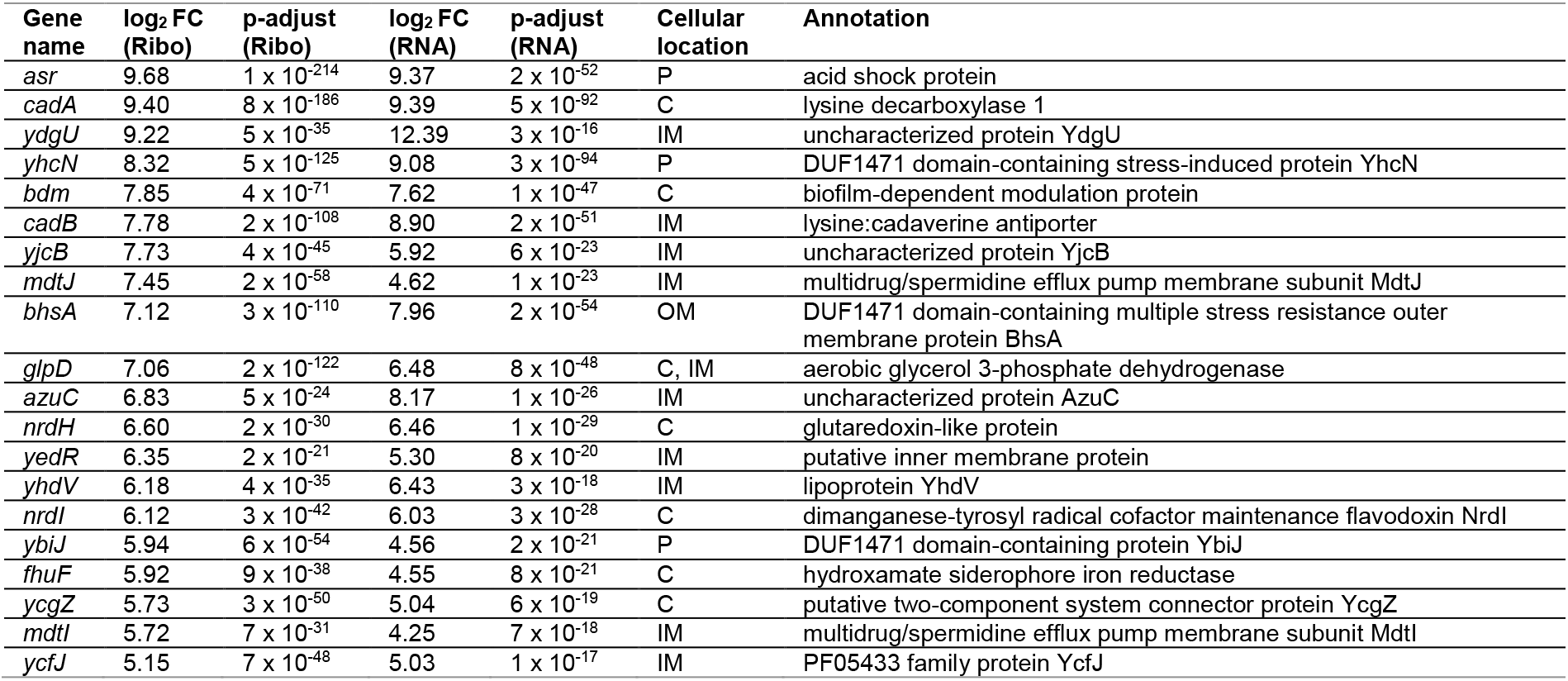
Top 20 genes with increased RPF levels at pH 4.4 compared to pH 7.6, sorted in descending order by Ribo-Seq log_2_ FC values. C, cytosol; P, periplasm; IM, inner membrane.

GSEA for biological processes identified the GO terms ‘enterobactin biosynthetic process’, ‘ferric-enterobactin import into cell’, ‘siderophore-dependent iron import into cell’ and ‘siderophore transmembrane transport’ as significantly enriched at pH 4.4 (Fig. 3). Specifically, the complete enterobactin biosynthesis pathway, comprising the *entCEBAH* operon, *entF*, *entH* and *ybdZ*, revealed significant enrichment under severe acidic conditions (Table S5). Furthermore, all subunits of the Ton complex (*tonB*, *exbB*, *exbD*) and its putative outer membrane receptor encoded by *yncD* exhibited significantly higher RPF and mRNA levels at pH 5.8 and pH 4.4 (Table S5). The Ton complex functions as a proton motive force-dependent molecular motor that facilitates the import of iron-bound siderophores.^68, 69^ Several other iron uptake systems, including a ferric dicitrate ABC transport system (*fecABCDE*), an iron (III) hydroxamate ABC transport system (*fhuACDB*), a ferric enterobactin ABC transport system (*fepA*, *fepB*, *fepCGD*) and a TonB-dependent iron-catecholate outer membrane transporter (*cirA*), were also induced under acidic conditions (Table S5). Moreover, the GO terms ‘protein maturation by iron-sulfur cluster assembly’ and ‘iron-sulfur cluster assembly’ were enriched at pH 4.4 (Fig. 3). Specifically, we detected a fivefold upregulation of all genes of the *isc* and *suf* operons (Table S5), which encode components of the complex machinery responsible for iron-sulfur cluster assembly in *E. coli*.^70^ In contrast, ‘heme transport’ was among the most downregulated biological process at both pH 5.8 and 4.4 (Fig. 3), which could potentially be the cause for iron limitation. Moreover, at low pH, the solubility of iron ions increases, which can destabilize iron-sulfur clusters.^71^ Iron limitation would be consistent with our data that *E. coli* upregulates the synthesis of iron-chelating siderophores and their transporters, as well as the components of the iron-sulfur assembly machinery. Given the better solubility of iron in a low pH environment, the question arises whether *E. coli* synthesizes siderophores to respond to iron limitation, or rather protects itself against an iron excess. The latter function has been demonstrated for *Pseudomonas aeruginosa,* where siderophores protected cells from harmful effects of reactive oxygen species. In this case, *P. aeruginosa* no longer secreted siderophores into the extracellular environment, and instead, stored them intracellularly.^72^ In conclusion, these results prompt the question whether the upregulation of the iron uptake machinery counteracts iron limitation or rather provides protection against iron excess under severe acid stress.

We also detected a significant enrichment for the GO terms ‘cellular response to acidic pH’, ‘stress response to copper ion’ and ‘copper ion transmembrane transport’ at pH 4.4 (Fig. 3). These results are in line with previous studies that have suggested an interplay between resistance to copper and acid stress in *Escherichia coli*.^73, 74^ This overlap between the two stress responses is further emphasized by our findings, because at pH 4.4, substantial upregulation of the Cu^+^ exporting ATPase CopA, and CusA, a component of the copper efflux system, was detected (Table S5). These results are of important physiological relevance, given that copper is an important antibacterial component in the innate immune system.^75, 76^

Among the downregulated genes at pH 4.4, the *tnaAB* operon and its leader peptide (*tnaC*), showed the most significant decrease in terms of RPF counts (Table 4). *tnaA* encodes a tryptophanase, which cleaves L-tryptophan into indole, pyruvate and NH_4_^+^, whereas *tnaB* encodes a tryptophan:H^+^ symporter.^77^ This finding is particularly intriguing because in a previous study persister cell formation in *E. coli* was related to lower cytoplasmic pH associated with tryptophan metabolism.^78^ It is important to note that we also detected a substantial upregulation in RPFs for *hipA* (Table S5), which encodes a serine/threonine kinase that plays a role in persistence in *E. coli*.^79^ Therefore, our data provide further evidence for the link between internal pH and persistence.

**Table 4:**
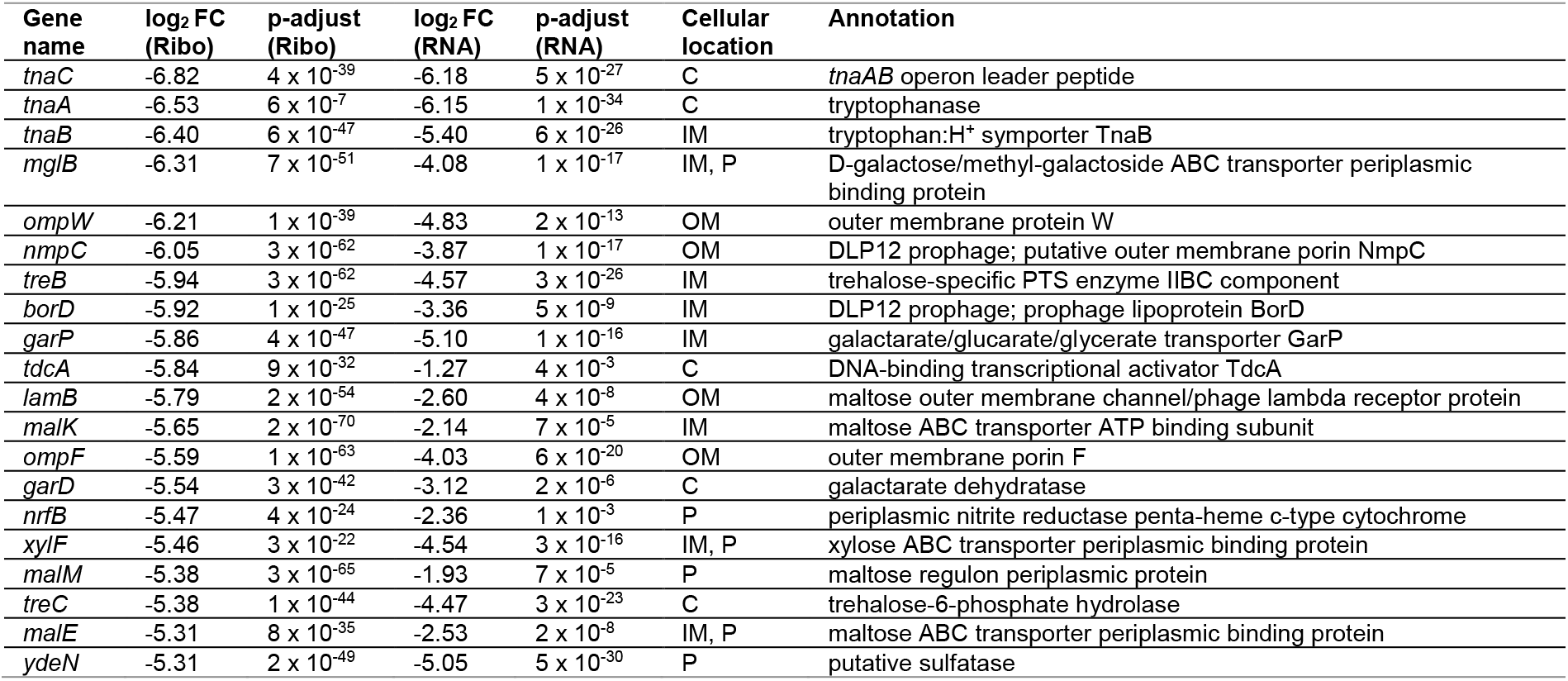
Top 20 genes with decreased RPF levels at pH 4.4 compared to pH 7.6, sorted in ascending order by Ribo-Seq log_2_ FC values. C, cytosol; P, periplasm; IM, inner membrane; OM, outer membrane.

The expression of several outer membrane proteins and porins (*ompW*, *ompF*, *nmpC*, *lamB*) was also downregulated at pH 4.4 (Table 4). This observation is consistent with the extensive restructuring of the *E. coli* lipid bilayers to reduce membrane permeability and limit proton entry. Similar to pH 5.8, the majority of the 20 proteins with the most reduced RPF levels compared with physiological pH are membrane proteins (Table 4). Moreover, the GO term ‘ATP synthesis coupled proton transport’ was significantly reduced at pH 4.4 (Fig. 3). This is explained by reduction of RPF levels of genes encoding subunits of the F_O_F_1_-ATPase (Table S5). The F_O_F_1_-ATPase uses the electrochemical gradient of protons to synthesize ATP from ADP and inorganic phosphate, but can also hydrolyze ATP to pump protons out of the cytoplasm.^80, 81^ As at pH 5.8, the most downregulated biological processes at pH 4.4 were almost exclusively GO terms related to transport and cellular metabolism (Fig. 3).

In summary, the response of *E. coli* to severe acid stress is dominated by the activation of survival strategies that limit the entry of protons into the cell, prevent protein aggregation, and maintain iron homeostasis. Severe acid stress leads to a reduction in metabolic, transcriptional and translational activity, thereby preparing *E. coli* for a dormant state. Eventually, these dormant cells may be able to withstand antibiotic attack (i.e. persister cells).

### 5. Expanding the regulatory network of enzyme-based H^+^-consuming acid resistance (AR) <u>systems</u>

Recently, we have shown that the Adi and Cad AR systems are mutually exclusively activated in individual *E. coli* cells, indicating functional diversification and division of labor under acid stress.^16^ Here, we analyzed the Ribo- and RNA-seq data to uncover further transcription factors (TF), which interconnect the enzyme-based H^+^-consuming AR2 to AR5 systems. The core components of the Gad system (AR2) (*gadA*, *gadB* and *gadC*) and several transcriptional components (*gadW*, *gadX*, *gadY*, *phoP*, *phoQ*) showed an increase in mRNA and RPF levels by ∼2-6-fold at pH 4.4, but not at pH 5.8, whereas the expression of *ydeO* was massively induced at pH 5.8 (particularly at the mRNA level) and RPF levels were decreased at pH 4.4 (Fig. S7). Expression of the core components of the Adi system (AR3), *adiA* and *adiC,* was induced at severe acid stress, but not at pH 5.8, consistent with our previous study.^16^ Upregulation was not detected for regulatory components of the Adi system. A novel finding was that the levels of *adiA,* but not *adiC,* were significantly higher in the Ribo-Seq data than in the RNA-Seq data (Fig. S7). In fact, *adiA* had the sixth highest increase in TE among all *E. coli* genes at pH 4.4 (Table S4), indicating translational regulation by a thus far unknown mechanism. The only other component of an AR system in *E. coli* known to be subject to translational regulation is the major regulator CadC of the Cad (AR4) system. CadC contains a polyproline motif and its translation therefore depends on the elongation factor P (EF-P), a process that keeps the copy number of CadC extremely low.^82^ As expected, expression of the core components of the Cad system (AR4), *cadA* and *cadB*, was tremendously increased at both pH 5.8 and 4.4. Genes of the Orn system (AR5) were not induced in our experimental setup (Fig. S7).

Next, we analyzed the mRNA and RPF levels of all annotated TFs to search for other potential TFs involved in the acid stress response of *E. coli* (Figs. 4A, 4B). At pH 5.8, YdeO showed by far the strongest induction at the transcriptional and translational levels, but for all other TFs, the expression levels hardly changed (Fig. 4A). At pH 4.4, the expression of numerous TFs was induced, including GadW, YdcI, and the antibiotic resistance-controlling regulator MarR. The strongest upregulation was found for the IclR-type regulator MhpR, and the iron-sulfur cluster-containing regulator IscR (Fig. 4B). Notably, while most acid-induced TFs were differentially expressed and displayed constant TE, YdcI exhibited constant mRNA levels but was differentially translated in response to acid stress (Fig. 4B). The contribution of all TFs with high FC values to survival under acid stress (Table S6) was tested in an acid shock assay. Cells of the corresponding knockout mutants^83^ and, for comparison, the *rcsB* and *gadE* mutants (each lacking a TF important for acid resistance) were exposed to pH 3 for 1 hour. All mutants except *marR* and *ydeO* showed significantly reduced survival compared to the parental strain (Fig. 4C). For *ydeO*, this result was consistent with our finding that transcript abundance and occupancy with ribosomes was upregulated at mild but not severe acid stress (Figs. 4, S7). Thus, YdeO appears to be only crucial under mild acid stress (Fig. 4A). In contrast, the *mhpR* mutant had a low survival rate comparable to that of *rcsB* and *gadE,* and the survival rates of the *iscR*, *ydcI,* and *gadW* mutants were only slightly higher (Fig. 4C). These results confirm the physiological relevance of these TFs for acid resistance.

**Figure 4:**
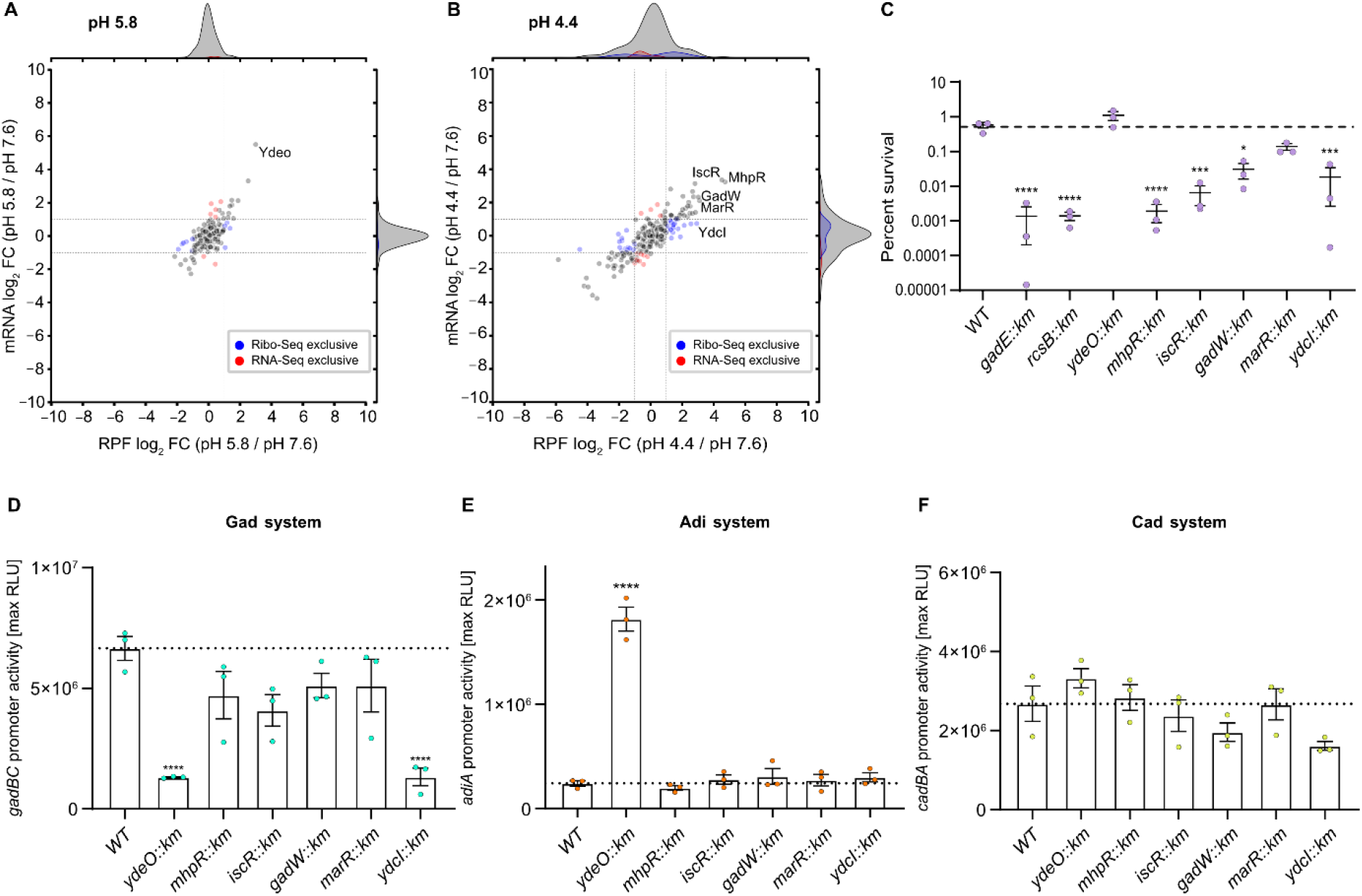
Contribution of transcription factors (TFs) to survival and AR induction under acid stress. Comparison of global RPF and mRNA log_2_ FC values of transcriptional regulators for **(A)** pH 5.8 vs. pH 7.6 and **(B)** pH 4.4 vs. pH 7.6. Dashed lines indicate log_2_ fold changes of +1 or −1. Changes detected exclusively by either RNA-Seq (red dots) or Ribo-Seq (blue dots) are colored. TFs described in panels C-F are highlighted. **(C)** Acid shock assay to test survival of *E. coli* BW25113 (WT) and the indicated mutants.^83^ Cells were grown in LB pH 7.6 to OD_600_ = 0.5. The cultures were split and then either grown at pH 7.6 or stepwise stressed (15 min pH 5.8, 15 min pH 4.4) before being exposed to LB pH 3 for 1 h. Colony forming units were counted and the ratio of surviving cells was calculated. The dashed line indicates the average percentage of surviving WT cells. **(D-F)** Luciferase-based promoter assays. WT and the indicated mutants were transformed either with plasmid pBBR1-P*gadBC:lux* **(D)**, pBBR1-P*adiA:lux* **(E)**, or pBBR1-P*cadBA:lux* **(F)** and grown in LB medium (pH 7.6) until OD_600_ = 0.5. The medium pH was then adjusted to 5.8 to induce the Cad system, or to pH 4.4 to induce the Adi and the Gad systems. Luminescence and growth were determined every 10 min in microtiter plates using a CLARIOstar Plus plate reader (BMG Labtech). Data are reported as relative light units (RLUs) in counts per second per OD_600_, with maximal RLU shown. Dashed lines indicate the average maximal RLU values of the WT. C-F: All experiments were performed in biological replicates (n = 3), and error bars represent standard deviations of the mean. ANOVA followed by Bonferroni’s multiple comparisons test was used to compare log-transformed max. RLU values between mutant strains and the wildtype (BW25113), (*p ≤ 0.05; **p ≤ 0.01, ***p ≤ 0.001, ****p ≤ 0.0001). FC, fold change; RPF, ribosome-protected-fragments.

Subsequently, we tested whether these TFs are involved in the regulation and interconnectivity of the Gad, Adi, and Cad systems. Therefore, we examined the promoter activities of *gadBC*, *adiA*, and *cadBA* in the corresponding knockout mutants^83^ using transcriptional reporter plasmids (promoter-*lux* fusions). The cultivation conditions were the same as those used for Ribo-Seq and RNA-Seq (Fig. 1A), and luciferase activity was monitored during growth in microtiter plates. We found that YdcI significantly affected the promotor activity of *gadBC* (Fig. 4D). Although the LysR-type regulator YdcI has been shown to affect pH stress regulation in *Salmonella enterica* serovar Typhimurium and *E. coli*, its precise role is still unclear.^84–86^ Based on the data presented here, we hypothesize that the decreased survival of the *ydcI* mutant under severe acid stress is due to decreased expression of the Gad system. The absence of YdeO resulted in an eightfold stimulation of the *adiA* promoter activity (Fig. 4E). Thus, YdeO not only activates the Gad system,^87^ but also appears to be a repressor for the Adi system. This implies that the Adi system is regulated not only by the XylS/AraC-type regulator AdiY, but also by YdeO. Thus, YdeO is the first example of a transcriptional activator shown to be involved in the regulation of more than one AR system in *E. coli* and might play a role in the heterogeneous activation of the Adi and Gad systems within a population. Although we observed a slight decrease in *cadBA* promoter activity in the *ydcI* mutant, the decrease was not statistically significant. Therefore, none of the tested TFs affected the Cad system (Figure 4F).

In conclusion, based on the differential expression data and lower survival of mutants during acid shock, we identified two novel TFs, namely MhpR and IscR, which are crucial under acid severe stress (Fig. 4C), but are not associated with the Gad, Adi and Cad systems (Figs. 4D – 4F). This implies that these regulators ensure survival of *E. coli* in acidic habitats by inducing other defense mechanisms. Of particular interest is MhpR, which had the highest increase in RPFs of all TFs at pH 4.4 (Fig. 4B), and the corresponding mutant had the lowest survival at pH 3 (Fig. 4C). Further studies are needed to determine whether MhpR, which is a specific regulator of the *mhpABCDFE* operon encoding enzymes for the degradation of phenylpropionate,^88, 89^ directly or indirectly contributes to acid resistance.

### 6. Differential expression of known and identification of novel sORFs under mild and severe acid stress

In recent years, the annotation of many bacterial genomes has been extended by previously unknown small proteins,^29^ many of which are located in the membrane.^90^ This progress has been achieved primarily through the development of optimized detection strategies using adapted ribosome profiling and mass spectrometry protocols.^28, 33, 91^ Recently, additional sORFs were identified in *E. coli* using antibiotic-assisted Ribo-Seq, which captures initiating ribosomes at start codons.^92, 93^ Advanced detection strategies also revealed novel small proteins in other species such as the archaeon *Haloferax volcanii*, the nitrogen-fixing plant symbiont *Sinorhizobium meliloti*, *Salmonella* Typhimurium and *Staphylococcus aureus*.^33–35, 94^

Among the previously known sORFs in *E. coli* K-12 and those discovered by Weaver *et al.* 2019,^92^ pH-dependent differential RPF levels were observed in our datasets for 12 and 29 small proteins at pH 5.8 and pH 4.4 respectively (Table S8). These findings validate the expression of these sORFs and highlight their physiological relevance in the acid stress response of *E. coli*. For example, induction of *mdtU*, an upstream ORF of *mdtJI*, was observed under mild acid stress (Fig. 5A) and corresponds to the observed upregulation of the multidrug/spermidine exporter MdtJI (Table 1). A previous study has shown that translation of MdtU is crucial for spermidine-mediated expression of the MdtJ subunit under spermidine supplementation at pH 9.^95^ A similar mechanism could operate under acid stress conditions. The strongest induction of sORFs under severe acid stress was detected for *ydgU* (located in the same transcriptional unit as the acid shock protein-encoding gene *asr*) and *azuC* (Fig. 5B). AzuCR acts as a dual-function RNA and encodes a 28-amino acid protein, but can also base pair as an sRNA (AzuR) with two target mRNAs including *cadA*.^96^ AzuCR modulates carbon metabolism through interactions with the aerobic glycerol-3-phosphate dehydrogenase GlpD.^96^

**Figure 5:**
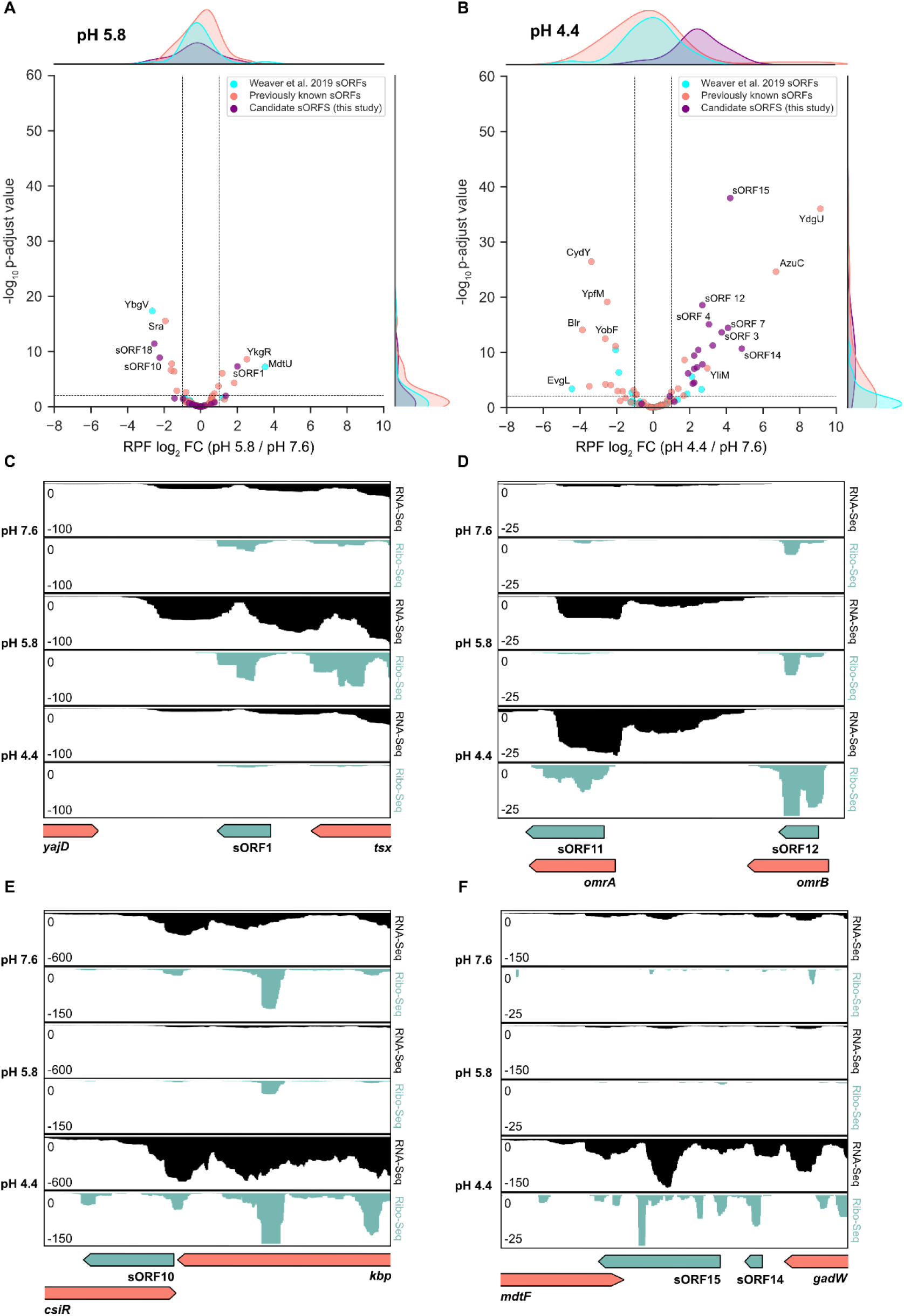
Differentially expressed sORFs upon exposure of *E. coli* to acid stress. Volcano plots illustrating differential RPF levels at **(A)** pH 5.8 vs. 7.6 and **(B)** pH 4.4 vs. 7.6 for sORF candidates identified in this study (Table S7), sORFs identified by Weaver *et al.* 2019^92^ and previously known sORFs. **(C-F)** JBrowse2 screenshots of read coverage from Ribo-Seq (green tracks) and RNA-Seq (black tracks) libraries at pH 7.6, pH 5.8 and pH 4.4. Schematic illustrations below indicate genomic locations of enriched sORFs under acidic stress and adjacent annotated genes. Novel sORF candidates sORF1 (13 aa), sORF10 (38 aa), sORF11 (28 aa), sORF12 (11 aa), sORF14 (13 aa) and sORF15 (93aa) are shown in green. aa - amino acids

In addition to known sORFs, we aimed to uncover further hidden small proteins on the basis that our Ribo-Seq data were acquired under stress conditions to which *E. coli* is exposed in its natural habitat, the gastrointestinal tract. In particular, we searched for novel sORFs that remained undetected in previous Ribo-Seq approaches when *E. coli* was grown at neutral pH.

Initial predictions for novel sORFs were acquired using the neural network-based prediction tool *DeepRibo*.^97^ All potential candidates were filtered based on coverage (rpkm >30 across all Ribo-Seq samples) and codon count [10 – 70 amino acids (aa)], with the exception of sORF15 (93-amino acids) (Table S7), which was manually discovered by inspecting the 3’UTR of *gadW*. To further refine our search, we focused on sORF candidates that were significantly induced at either pH 5.8 or pH 4.4 (RPF log_2_FC > 2 and p-adjust < 0.05) compared to pH 7.6. Predictions that overlapped with annotated genes on the same strand were excluded because Ribo-Seq signals were indistinguishable. This workflow yielded 152 candidates that were visually inspected using the Web-based genome browser JBrowse2.^98^ Candidates with continuous coverage across the predicted sORF, matching the ORF boundaries, and promising Shine-Dalgarno sequences were considered high-confidence candidates. In total, we identified 18 acid-induced sORF candidates (Table S7) that had not been previously detected. Of note, most of the candidates are encoded as part of operons or are located in the 3’UTR of annotated genes. In addition, we detected one independent antisense sORF (sORF2 encoded antisense to *tesA*) and two upstream ORFs (leader peptides): sORF18, located upstream of the translation start site of the periplasmic chaperone encoding *osmY*, and sORF8, located close to the glucokinase-encoding gene *glk* (Table S7).

Of these 18 acid-induced candidate small proteins (Table S7), 17 had higher RPF counts at pH 4.4 compared to pH 5.8. This suggests that the contribution of sORFs to acid defense in *E. coli* is more relevant under severe acid stress. Only sORF1 showed a higher expression level in cells exposed to mild acid stress (Fig. 5C). sORF1 is located in the 3’UTR of *tsx*, which encodes a nucleoside-specific channel-forming protein. This finding is consistent with the observed increased requirement for nucleotides by *E. coli* at pH 5.8 (Fig. 3).

For the first time, we identified two sORF candidates located within genes encoding the redundant small regulatory RNAs OmrA and OmrB (Fig. 5D). *omrA* and *omrB* are highly identical at the 3’ and 5’ ends, differ mainly in their central parts, and regulate the expression of numerous outer membrane proteins.^99^ Our analysis suggests that both OmrA and OmrB act as dual-function RNAs under severe acid stress and encode small proteins: a 28-amino acid protein OmrA (sORF11) and an 11-amino acid protein OmrB (sORF12) (Fig. 5D). Due to the sequence variation in the central parts, translation of OmrB ends at an earlier stop codon. Notably, both *omrA* and *omrB* displayed higher RPF levels at pH 4.4, whereas transcription of *omrA* but not *omrB* was induced at pH 4.4 (Fig. 5D). Thus, despite the high sequence similarity, *omrA* and *omrB* do not encode identical small proteins under severe acid stress and are differentially regulated at the transcriptional and translational levels. We also detected an acid-induced sORF (sORF3) in *rybB* (Table S7), another sRNA involved in the regulation of outer membrane proteins.^100^ To our best knowledge, the presence of OmrA, OmrB and RybB peptides has not yet been reported.

Three new candidate sORFs potentially involved in the regulation of AR systems (see chapter 5) were detected. sORF10 is located in the 3’ UTR of a potassium binding protein encoded by *kbp*, and encoded antisense to the transcriptional regulator CsiR (Fig. 5E). The latter might be involved in the regulation of the Adi system.^16, 101^ Given the significant upregulation of sORF10 at pH 4.4 and its complete complementarity to the 3’end of the *csiR* mRNA, we hypothesize that sORF10 plays a role in fine-tuning the expression of the Adi system. Strikingly, we also discovered two high-confidence candidates for sORFs located in the relatively long 3’UTR of GadW, one of the major transcriptional regulators of the Gad system (Fig. 5F). sORF14 and sORF15 exhibit constant coverage across the predicted ORF and contain Shine-Dalgarno sequences (Table S7). These results suggest that the complex Gad system may consist of even more components.

To gain further insight into the subcellular location and features of the newly identified sORF candidates, we used PSORTb^102^ and DeepTMHMM^103^ for transmembrane topology prediction. Notably, sORF15 is predicted to be located in the inner membrane and has two transmembrane helices which were predicted with a probability of > 90% (Fig. S8A). Additionally, the sORF15 protein structure prediction, using AlphaFold2^104^ in Google Colab (ColabFold),^105^ revealed a potential third helix towards the C-terminal end (Fig. S8B). Using blastp and tblastn,^106^ we found homologs of sORF15 with > 80% identity in *Vibrio*, *Shigella*, *Klebsiella*, *Salmonella*, *Enterococcus* and *Escherichia* (Fig. S8C) and identified homologs with at least 60 % identity for approximately half of the other candidate sORFs (sORF2, 4, 5, 6, 7, 9, 10, 11, 16 and 18). These results strengthen confidence in the correct prediction of these sORFs. However, homologs in other species often only displayed partial matches and were almost exclusively annotated as ‘hypothetical proteins’, as illustrated for sORF15 (Figure S8C).

In conclusion, we identified 18 high-confidence candidates for novel sORFs that are significantly induced upon exposure of *E. coli* to mild or severe acid stress.

### 7. Differentiation of the acid stress and general stress response using autoencoder-based machine learning

In general, stress response mechanisms can be broadly classified into two categories: global stress responses and adaptations to specific types of stress. Global stress responses can be triggered by various stimuli and provide protection against multiple other unrelated stress factors.^107^ The global response often involves the activation of alternative sigma factors that affect hundreds of genes. In contrast, adaptations to specific types of stress are tailored to the specific stressor and involve a regulator that senses an environmental cue and modulates the expression of a set of genes, which counteract the stress.^107, 108^

Given the large number of differentially regulated genes and pathways in response to acid stress (Figs. 2, 3), we asked which of these adaptive mechanisms are acid-specific and which are also triggered by other stressors. In order to distinguish acid-specific and general stress responses, we used denoising autoencoders (DAEs): deep learning models designed for meaningful dimensionality reduction.^109, 110^ DAEs accomplish this by passing data through an encoder that compresses it into activations of a bottleneck layer (Fig. 6A1), with each node in the bottleneck layer interpretable as a coordinated expression program.^111^ For our analysis, we employed an ensemble of deep DAEs (see Methods),^111^ trained on the *E. coli* K-12 PRECISE 2.0 compendium^112^ augmented with additional stress conditions^113^ as well as the acid stress conditions of the current study (Fig. 6A1). Using this method and data set, we have conducted a comparative analysis of the transcriptional response of *E. coli* to pH 4.4 and pH 5.8, contrasted against an extensive range of other stress conditions including heat stress,^114^ ethanol stress,^115^ osmotic stress,^116^ oxidative stress,^117, 118^ low oxygen (LOX)^113^ and exposure to sublethal concentrations of chloramphenicol (CAM)^113^ and trimethoprim (TMP).^113^

**Figure 6:**
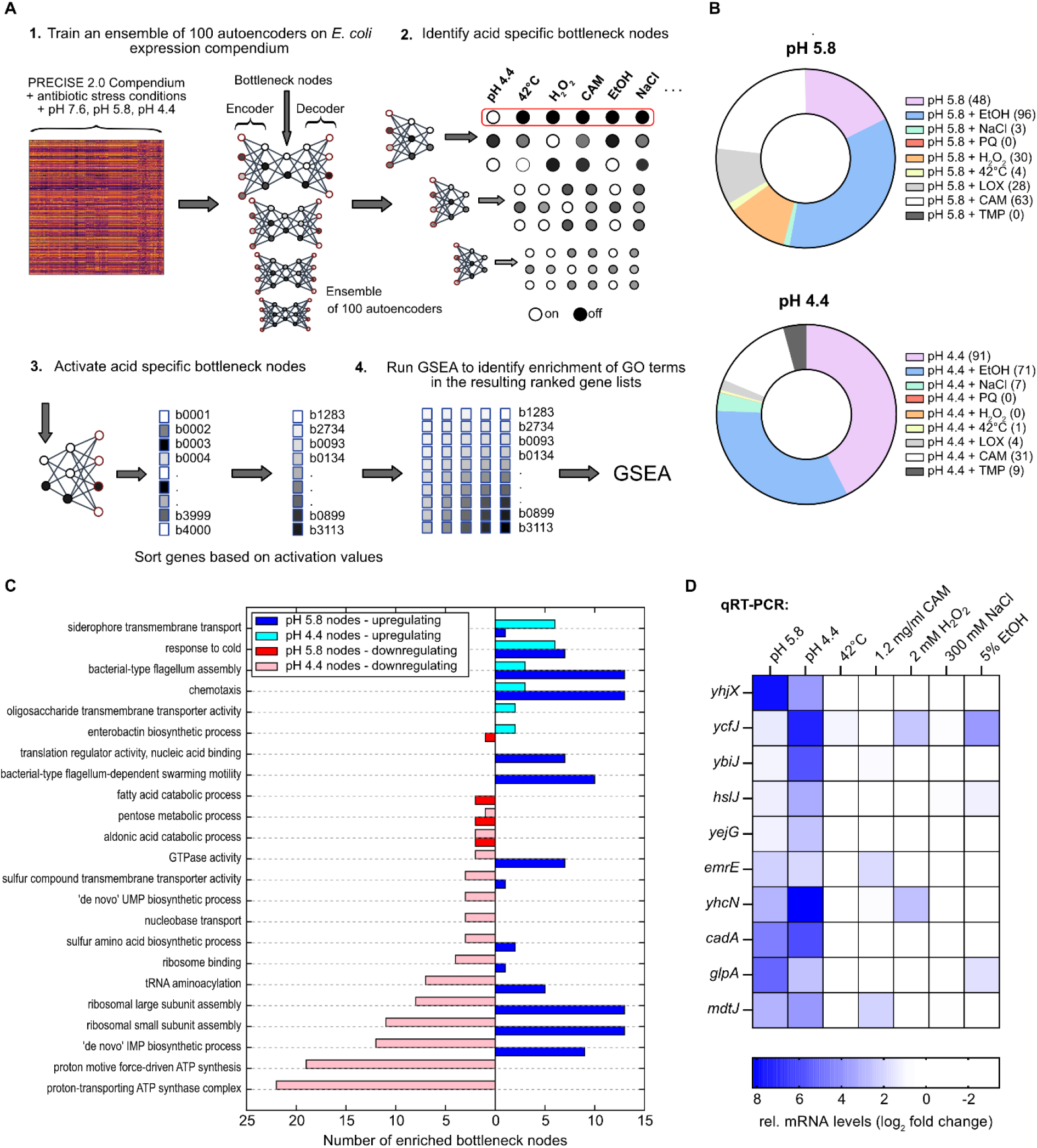
Differentiation between acid stress response and general stress using autoencoders. **(A)** Schematic overview of the autoencoder ensemble training and subsequent bottleneck group identification pipeline. **(B**) Donut charts indicating the proportion of bottleneck nodes which turn on only under the specified conditions. Absolute numbers of specific bottleneck nodes are listed in brackets after each condition. **(C)** Significantly enriched GO terms associated with pH 4.4 and pH 5.8 specific nodes. Left-facing red bars indicate nodes which downregulate the corresponding GO term, while right-facing blue bars indicate upregulation. **(D)** Verification of acid stress specific genes predicted by autoencoders. *E. coli* cells were either grown as indicated in Figure 1A, or exposed to oxidative (H_2_O_2_), osmotic (NaCl), antibiotic (chloramphenicol), or heat (42°C) stress. Total RNA was isolated, and relative mRNA levels were measured by RT-qPCR. Fold change values were determined relative to non-stress conditions and normalized using either *secA* or *recA* as reference genes. Standard deviations were calculated from three replicates (n =3) and accounted for < 10% of fold change values in all cases. EtOH = ethanol, PQ = paraquat, LOX = low oxygen, CAM = chloramphenicol, TMP = trimethoprim.

To identify biological processes associated with a particular stress condition, we passed the associated RNA-seq data set into the encoder of each network and identified bottleneck nodes which uniquely turn on by that data set (Fig. 6A2). We then manually turned on these nodes to generate gene sets which are associated with that condition, which can be further analyzed through GO term enrichment (Fig. 6A3,4). Using this procedure, we identified groups of nodes which uniquely turn on for acid stress conditions and turn off in all other stress conditions, as well as groups which are simultaneously on for both acid and one additional stress condition. We observed that there are many nodes which turn simultaneously on upon both acid and ethanol exposure (Fig. 6B). The overlap between acid stress and ethanol stress responses has been noted previously and can be explained by the fact that ethanol fluidizes the cytosolic membrane and increases the permeability for protons.^119^ Further, there are indications of an overlap between acid and antibiotic stress,^120, 121^ reflected in the high number of acid + CAM activating nodes (Fig. 6B).

To pinpoint which cellular adaptations cause acid-specificity for the 48 and 91 specific bottleneck nodes at pH 5.8 and 4.4 (Fig. 6B), respectively, we conducted GSEA for biological processes on each of the gene sets associated with acid specific node groups (see Methods section). The GO terms that were significantly enriched in the highest number of both pH 5.8- and pH 4.4-specific upregulating node gene sets were ‘siderophore transmembrane transport’, ‘response to cold’, ‘bacterial-type flagellum assembly’ and ‘chemotaxis’ (Fig. 6C), reflecting our previous differential RNA-seq analysis (Figure 3). The appearance of the GO term ‘response to cold’ might be a result of the lack of cold stress in our compendium of stressors. Additionally, it should be noted that genes associated to this GO term include cold shock proteins which may have broader roles in survival of stress conditions,^122, 123^ as well as several prophage genes and ribosome biogenesis factors. We found that mild acid stress turns on nodes which correspond to gene sets associated with nucleotide and ribosome biosynthesis, including the GO terms ‘ribosome large subunit assembly’, ribosome small subunit assembly’ and ‘*de novo* IMP biosynthetic process’, while severe acid stress turns them off (Fig. 6C). These findings are consistent with our previous observations (Chapters 3 and 4), namely that *E. coli* induces nucleotide and ribosome biosynthesis to cope with mild acid stress, but enters a metabolically inactive state under severe acid stress. The GO terms significantly affected in the highest number of acid-specific pH 4.4 downregulated bottleneck nodes were ‘proton motive force-driven ATP synthesis’ and ‘proton-transporting ATP synthase complex’ (Fig. 6C). These two GO terms exclusively involve genes encoding subunits of the F_O_F_1_ ATP synthase and can be considered paradigms for acid-specific adaptations since the F_O_F_1_ ATP synthase can also pump protons.^124^

Considering that we detected a high number of genes induced by severe acid stress with unknown functions (Table 3) and lacking GO associations, we expanded our search for acid-specific adaptations from GO terms to single genes. In order to select acid-specific candidate genes, we investigated all genes associated to acid-specific bottleneck nodes and calculated the log_2_FC between each acid stress and every other above-mentioned stress condition for these genes. Genes with the highest expression values under acidic conditions and log_2_FCs of at least 0.5 for at least 95% of comparisons were then selected. This procedure yielded 10 candidate genes (evaluated in Figure 6D). To experimentally validate that these genes are indeed specifically upregulated under acid stress, we exposed *E. coli* to a variety of common stressors and performed qRT-PCR. *E. coli* cells were either grown under acid stress (Figure 1A), or exposed to oxidative (H_2_O_2_), osmotic (NaCl), antibiotic (chloramphenicol), or heat (42°C) stress. For all investigated genes, the strongest upregulation was observed at either pH 4.4 or pH 5.8 relative to non-stress conditions (Fig. 6D), except for *ycfJ* which was activated at pH 4.4 and under oxidative and ethanol stress. The remaining investigated genes were only upregulated under one other stress condition at most (Fig. 6D). Given that *emrE* and *mdtJ* encode multidrug exporters, the induction upon supplementation with sublethal concentrations of chloramphenicol is not surprising and further underscores the interplay between acid and antibiotic stress. The observed upregulation of *yhcN* under oxidative stress (Fig. 6D) was also reported previously.^125^ Nevertheless, we uncovered four *bona fide* examples of genes (*ybiJ*, *hslJ*, *yejG* and *yhjX*) that displayed exclusive pH-dependent expression (Fig. 6D). Induction of *yhjX,* encoding a putative pyruvate transporter, might be related to the deamination of serine, which yields ammonia and pyruvate in uropathogenic *E. coli*.^126^ The precise molecular functions of YbiJ, YejG and HslJ in the context of acid stress are currently unclear. These results highlight that our autoencoder pipeline is complementary to differential gene expression analysis, yielding biologically consistent results while also identifying expression patterns that uniquely discriminate acid stress from other stress responses.

## Conclusions

Here, we present the first comprehensive study on the global transcriptome- and translatome-wide response of *E. coli* exposed to varying degrees of acid stress. Our investigation goes beyond previous research which focused on comparing *E. coli* transcriptomes across different pH levels during growth.^18, 21^ Instead, we report on rapid changes occurring upon sudden pH shifts, which are relevant for bacteria, such as *E. coli*, during passage of the gastrointestinal tract.^127^

Using both Ribo- and RNA-Seq, we uncovered not only well-known acid defense mechanisms, but also numerous previously undiscovered relevant genes and pathways to combat mild and severe acid stress (Fig. 7). The latter include, siderophore production, glycerol-3-phosphate conversion, copper export, *de novo* nucleotide biosynthesis and spermidine/multidrug export (Figs. 3, 7). A striking number of membrane proteins and H^+^-coupled transporters were found to be downregulated under both mild and severe acid stress (Fig. 7, Tables 2, 4), underscoring the importance of the cytosolic membrane and its composition as a barrier for protons. Moreover, under severe stress, many outer membrane proteins were downregulated (Fig. 7, Table 4). Notably, a large proportion of genes with yet unknown functions was strongly induced, particularly under severe acid stress (Table 3). Our approach implies that exposing *E. coli* to culture conditions mimicking near-lethal habitats can offer valuable insights into the molecular functions of genes with low expression levels under standard growth conditions.

**Figure 7:**
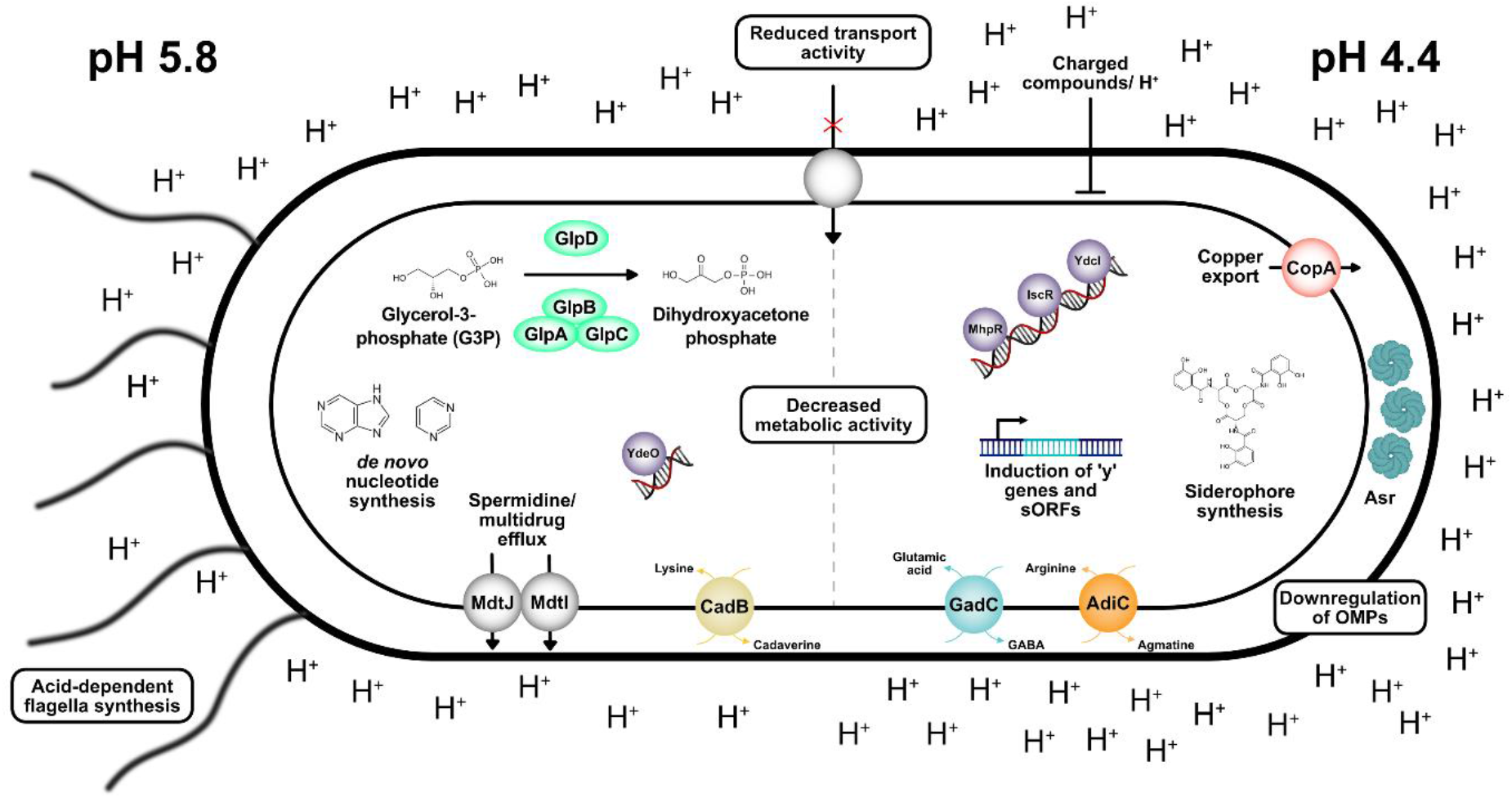
Overview of the fine-tuned response of *E. coli* to mild (pH 5.8) and severe (pH 4.4) acid stress. Acid stress counteracting mechanisms including those revealed by Ribo-Seq and RNA-seq are indicated.

Our analysis revealed two new TFs, MhpR and IscR, involved in acid stress adaption. Furthermore, we gained new insights into the role of the TFs YdeO and MarR. YdeO controls not only transcription of genes of the Gad system but also *adiA* of the Adi system (Fig. 4), suggesting that YdeO connects the regulation of two AR systems in *E. coli*. The observed upregulation of MarR under acid stress but the low contribution of this TF to acid resistance (Fig. 4B) may provide a link to antibiotic resistance and solvent stress tolerance in *E. coli*.^128^

In addition to the pH-dependent differential expression levels of previously identified small proteins, such as YdgU, MdtU and AzuC, we identified 18 high-confidence not yet annotated sORF candidates (Fig. 5). Of particular interest are sORF14 (13-amino acids) and sORF15 (93-amino acids), which are located in a transcriptional unit with *gadW* and *gadX,* suggesting their association with the Gad AR system and a potential involvement in glutamate transport and/or glutamate decarboxylation to gamma-aminobutyrate (GABA). Considering the predicted membrane location of sORF15 and its adjacent gene *mdtF*, an association with either the glutamate/GABA antiporter GadC and/or the multidrug efflux pump MdtF is conceivable.

The autoencoder-based comparison with other common stressors allowed us to distinguish acid stress-specific adaptations from general stress response programs (Fig. 6). Thereby it was possible to differentiate between direct and indirect effects triggered by protonation and/or cellular damage. Considering the growing volume of next generation sequence data, denoising autoencoders will be an increasingly important tool for interpreting future studies in the full context of accumulating RNA-seq data sets. Colonizing the intestinal tract is a complex process that does not only include rapid pH changes, but also alterations in oxygen and nutrient availabilities as well as competition with other bacteria. The ability of pathogenic *E. coli* strains to respond to such rapidly changing environments ensures their fitness advantage. We have shown here, for acid stress, the complexity of the regulatory network for ensuring survival and adaptation. The use of autoencoders, successfully tested here, could allow for identification of physiological weak points associated with survival of specific stresses. Targeting such weak points could lead to new classes of antibiotics or antivirulence treatments that take advantage of the unique expression patterns induced by natural stress conditions encountered in the host environment.

## Supporting information

Key resources table

## Author contributions

Conceptualization, K.S. and K.J.; Methodology, K.S., R.G., W.K.C., L.B., R.B and K.J.; Investigation, K.S., R.G and W.K.C.; Software, R.G., W.K.C., L.B. and R.B.; Writing – Original Draft, K.S. and K.J.; Writing – Review & Editing, K.S., R.G., W.K.C., L.B., R.B. and K.J.; Funding Acquisition and Resources, L.B., R.B. and K.J.; Supervision, K.J.

## Acknowledgement

We thank Stefan Krebs and Helmut Blum for conducting Illumina sequencing and acknowledge Christoph Kurat for providing the Freezer mill machine. We thank Sophie Brameyer for scientific discussions, Pol Bannasch and Annika Krimmel for experimental assistance, and Lydia Hadjeras for help with sORF detection. In addition, K.S. acknowledges the support of the Graduate School Life Science Munich (LSM). This work was financially supported by the Deutsche Forschungsgemeinschaft (DFG, German Research Foundation): Project number 471254198-JU270/ 22-1, Project number 395357507 – SFB 1371 to K.J. and grant BA 2168/21–2 to R.B.; and by the Bavarian State Ministry for Science and the Arts through the research network bayresq.net to L.B. Computational resources were provided by the BMBF-funded de.NBI Cloud within the German Network for Bioinformatics Infrastructure (de.NBI) (031A532B, 031A533A, 031A533B, 031A534A, 031A535A, 031A537A, 031A537B, 031A537C, 031A537D, 031A538A).

## METHODS

### RESOURCE AVAILABILITY

#### Lead contact

Further information and requests for resources and reagents should be directed to and will be fulfilled by the lead contact, Kirsten Jung.

#### Materials availability

New plasmids generated in this study are available upon request via the lead contact.

#### Data and code availability

The raw sequencing data for both the Ribo-seq and RNA-seq was deposited at Gene Expression Omnibus (GEO) under the accession GSE219022.

A JBrowse2^98^ online genome browser instance containing annotation files and read coverage files of the processed data is available via the RIBOBASE (http://www.bioinf.uni-freiburg.de/ribobase).

Additional information required to reanalyze the data reported in this paper is available from the lead contact upon request.

### EXPERIMENTAL MODEL AND SUBJECT DETAILS

Bacterial strains and plasmids used in this study are listed in Table S9. *E. coli* strains were cultivated in lysogeny broth (LB) medium (10g/l tryptone, 5 g/l yeast extract, 10 g/l NaCl) and incubated aerobically in a rotary shaker at 37°C. When appropriate, media were supplemented with 15 µg/ml gentamycin.

For ribosome profiling and RNA-Seq experiments, the pH of the medium was adjusted at the indicated time points by direct addition of 5 M HCl to the growing cultures (Fig. 1A).

For comparison with other stress conditions (heat stress, osmotic stress, oxidative stress antibiotic stress and ethanol stress), *E. coli* was initially grown in LB medium to OD_600_ = 0.5. Stress conditions were initiated either by moving flasks to a pre-heated 42°C incubator, or by addition of either H_2_O_2_ [2 mM final concentration (f.c.)], NaCl (300 mM f.c.), chloramphenicol (1.2 µg/ml f.c.), or ethanol [5% (v/v) f.c.]. In all cases, samples were collected after 15 min of stress treatment.

### METHOD DETAILS

#### Plasmid construction

Molecular methods were performed according to standard protocols or according to the manufactureŕs instructions. Kits for isolation of plasmids and the purification of PCR products were purchased from Süd-Laborbedarf (SLG). Enzymes and HiFi DNA Assembly Master Mix were purchased from New England Biolabs. To construct the reporter plasmid pBBR1-MCS5-P*_gadBC:lux_*, 335 nt of the upstream region of *gadBC* were amplified by PCR using primers KSO-0131/KSO-0132 (Table S10) and MG1655 genomic DNA as a template. After purification, the promoter fragment was assembled into a PCR linearized pBBR1-MCS5 plasmid via Gibson assembly.^129^ Correct insertion was verified by colony PCR and sequencing.

#### Sample collection for Ribo-Seq and RNA-Seq

Three sets of biological triplicates of MG1655 cells were inoculated to a starting OD_600_ of 0.05 from overnight cultures and grown to exponential phase (OD_600_ = 0.5) in 200 ml of unbuffered LB medium (pH 7.6). Two sets of cultures were adjusted to pH 5.8 by direct addition 5 M HCl, while one set was further grown for 30 min at pH 7.6 (Fig. 1A). After 15 min, one set of biological triplicates was further adjusted from pH 5.8 to pH 4.4 by addition of 5 M HCl, while the other cultures remained at pH 5.8 or pH 7.6, respectively (Fig. 1A). Cells were grown for another 15 min, samples were collected, and Ribo-Seq and RNA-Seq was performed as described below. pH values before and after pH-shifts, as well as final optical densities were monitored throughout the experiment (Tables S1 and S2).

#### Ribosome profiling

Whole culture flash freezing, cell lysis using a freezer mill, pelleting ribosomes over sucrose cushions and ribosomal footprint isolation using a size selection gel were performed following the published protocol from Mohammad and Buskirk.^39^ MNase treatment, monosome recovery, RNA isolation, end-labelling by T4 Polynucleotide Kinase and cDNA library construction were conducted by adapting the protocol from Latif and colleagues.^38^

Briefly, 100 ml of liquid cultures and 10x lysis buffer were flash frozen in liquid nitrogen. For each sample, 90 g of frozen cells were mixed with 10 g of frozen 10x lysis buffer and lysed by cryogenic grinding in a SPEX SamplePrep 6875 Freezer/Mill (10 cycles, 10 Hz, 5 min precool, 1 min run, 1 min cool). The pulverized samples were thawed and the lysate was pre-cleared by centrifugation (9,800 *g*, 10 min, 4°C) in a Beckman Coulter Optima XE-90 Ultracentrifuge using a 50.2 Ti Rotor. The supernatant was used to pellet ribosomes over sucrose cushions by centrifugation in a Beckman Coulter Optima XE-90 Ultracentrifuge using a 70.1 Ti Rotor (330,000 *g*, 1.5 h, 4°C). After resuspension of pellets, nuclease digestion was performed using MNase (NEB) (2 h, 25°C). Monosomes were recovered using MicroSpin S-400 HR columns (GE Healthcare) and RNA was isolated using the miRNeasy Mini Kit (QIAGEN) in combination with the RNase-Free DNase Set (QIAGEN). The isolated RNA was loaded on a 15% TBE Urea size selection gel and after staining with SYBR Gold (Invitrogen), ribosomal footprints between 15 and 45 nt were excised from the gel. For elution of RNA, gel pieces were crushed by poking a hole with an 18 G needle in a 0.5 ml tube, placing it in a 1.5 ml tube and subsequent centrifugation. RNA was recovered by precipitation after adding elution buffer to the crushed gel, overnight incubation (4°C) and centrifugation in Corning-Costar Spin-X Centrifuge Tube Filters (20,000 *g*, 3 min, room temperature). The isolation of RNA fragments corresponding to ribosomal footprints of 15 – 45 nt size was verified using the RNA 6000 Nano Kit (Agilent) and the Agilent 2100 Bioanalyzer. 5’ phosphorylation of RNA fragments was achieved using T4 Polynucleotide Kinase (NEB) and Adenosine 5’-Triphosphate (ATP) (NEB). After RNA recovery using an RNA MinElute Cleanup Kit (QIAGEN), footprint fragments were again evaluated using the Agilent 2100 Bioanalyzer as described above. cDNA libraries were constructed using the NEBNext Small RNA Library Prep Set for Illumina (NEB) with 14 PCR amplification cycles. cDNA library quality was assessed using a High Sensitivity DNA Kit (Agilent). cDNA libraries were purified using the QIAquick PCR Purification Kit (QIAGEN) and sequenced using a HiSeq 1500 machine (Illumina) in single-read mode with 50 bp read length.

#### RNA-Seq analysis

After flash-freezing the cultures for Ribo-Seq, 6 ml of the remaining culture volume were mixed with 1.2 ml Stop Mix solution [95% (vol/vol) ethanol, 5% (vol/vol) phenol] to terminate ongoing transcription and translation. Samples were frozen in liquid nitrogen and stored (−80°C) until RNA isolation. Cells were pelleted (3,000 *g*, 15 min, 4°C) and total RNA was isolated using the miRNeasy Mini Kit (QIAGEN) in combination with the RNase-Free DNase Set (QIAGEN). Integrity of RNA samples was evaluated using the RNA 6000 Nano Kit (Agilent) and the Agilent 2100 Bioanalyzer. RNA was quantified using the Qubit RNA HS Assay Kit (Invitrogen). Ribosomal RNA depletion was performed using the NEBNext rRNA Depletion Kit for bacteria (NEB) and directional cDNA libraries were prepared using the NEBNext Ultra II Directional RNA Library Prep Kit for Illumina (NEB). cDNA library quality was assessed using a High Sensitivity DNA Kit (Agilent). Finally, cDNA libraries were sequenced using a NextSeq 1000 machine (Illumina) in single-read mode with 60 bp read length.

#### Next generation sequencing data analysis

*E. coli* Ribo-seq and RNA-seq raw sequencing libraries were processed and analyzed using the published HRIBO workflow (version 1.6.0),^40^ which has previously been used for analysis of bacterial Ribo-seq data.^35^ HRIBO is a snakemake^130^ workflow that downloads all required tools from bioconda^131^ and singularity.^132^ All necessary processing steps are automatically determined by the workflow.

Adapter and quality trimming of the reads was performed using cutadapt (version 2.1).^133^ The trimmed reads were then mapped against the *E. coli* K-12 MG1655 reference genome (ASM584v2) with segemehl (version 0.3.4).^134^ Reads corresponding to ribosomal RNA (rRNA), transfer RNA (tRNA) and multi-mapping reads were removed with SAMtools (version 1.9)^135^ using rRNA and tRNA annotations.

Quality control was performed by creating read count statistics for each processing step and RNA-class with Subread featureCounts (1.6.3).^136^ All processing steps were analyzed with FastQC (version 0.11.8)^137^ and results were aggregated with MultiQC (version 1.7).^138^ Additionally, a principle component analysis (PCA) was performed to determine whether the major source of variance in the data stems from the different experimental conditions. The PCA ensures correctness of the downstream differential expression analysis. To generate the PCA plots, normalized read counts for all samples were generated using DESeq2^139^ normalization. To improve the clustering, a regularized log transform was applied to the normalized read counts. Standard deviations and variance were subsequently calculated using DESeq2 and the first three principle components were plotted using plotly.^140^

Read coverage files were generated with HRIBO using different full read mapping approaches (global or centered) and single-nucleotide mapping strategies (5’ or 3’ end). Read coverage files were normalized using the counts per million (mil) normalization. For the mil normalization, read counts were normalized by the total number of mapped reads within the sample and scaled by a per-million factor.

Metagene analysis of ribosome density at start codons was performed as described previously.^141^ Here, annotated start codons of coding sequences are collected and the density of the read coverage is determined for every position in a pre-determined window around the collected start codons.

A differential expression and translation analysis was performed using the tool deltaTE.^43^ The tool combines both Ribo-seq and RNA-seq data by calculating the translational efficiency (TE) of genes in order to capture changes in translational regulation when comparing different growth conditions.

#### Gene set enrichment analysis

Gene Set Enrichment Analysis (GSEA) was conducted using the tool *clusterProfiler.*^51^ To this end, the genome wide annotations database for *E.coli* K-12 MG1655^142^ was used in combination with the results of the differential expression analysis. The analysis was focused on the biological process (BP) domain, omitting the cellular component (CC) and molecular function (MF) domains. Further, the minimum gene set size was set to 5 and the maximum to 30. To avoid redundancy within the results, GO terms were filtered and only terms at the bottom (lowest branch level) of the GO hierarchy were analyzed.

#### Detection and differential expression of novel acid-induced sORF candidates

sORF candidates were initially detected using the neural network *DeepRibo*^97^. Summary statistics including TE, rpkm, codon counts, nucleotide and amino acid sequences, for annotated and potential novel sORFs were computed using HRIBO (version 1.6.0).^40^ Moreover, GFF track files were created for detailed manual inspection using the web-based genome browser JBrowse2.^98^

*DeepRibo* was reported to produce high numbers of false positives.^143^ Therefore, the high number of initial predictions (> 25.000) was reduced by introducing cut-off criteria based on sORF length and Ribo-Seq coverage. Predictions with average rpkm (reads per kilobase of transcript per million mapped reads) values < 30 across all Ribo-Seq samples and outside of the codon count range of 10 – 70 amino acids were excluded from further analysis. To specifically detect sORFs, which are only detectable under acidic conditions and involved in the acid response of *E. coli*, the search was further restricted to predictions with deltaTE Ribo- and RNA log_2_FC values of ≥ 2, in combination with p-adjust values of ≤ 0.05, at either pH 5.8, or pH 4.4, compared to pH 7.6. Additionally, *DeepRibo* predictions overlapping annotated genes on the same strand were also excluded. The remaining 152 candidates were manually inspected in JBrowse2, and novel sORFs were included in our final candidate list (Table S7) if the coverage was even over the predicted ORF, restricted to the ORF boundaries, and potential Shine-Dalgarno sequences were detectable. The remaining candidates were manually curated and, in a few cases, alternative sORFs in the genomic vicinity of DeepRibo predictions were selected, which matched better to the Ribo-Seq coverage signal. Differential expression values, comparing pH 7.6, pH 5.8 and pH 4.4, for novel sORF candidates, previously known sORFs, and sORFs detected by Weaver *et al.* 2019^92^ were calculated using *deltaTE*.^43^

Homologs for novel sORFs were identified using blastp and tblastn^106^ within the ‘BLAST at NCBI’ function in the CLC Main Workbench v. 20.0.4 (QIAGEN). For tblastn, the following parameters were used: an Evalue (Expect value) ≤ 0,05, a seed length that initiates an alignment (word size) of 6, and the filter for low complexity regions off. Potential cellular localization was evaluated with PSORTb v3.0.3 (https://www.psort.org/psortb/)^102^ and transmembrane topology was assessed using DeepTMHMM v1.0.24.^103^ The protein structure of sORF15 was predicted by running AlphaFold2^104^ via Google Colab (ColabFold v1.5.2)^105^ with default settings.

#### RNA isolation and RT-qPCR analysis

RNA was isolated using the Quick-RNA Miniprep Kit (Zymo Research), or miRNeasy Mini Kit (QIAGEN) in combination with the RNase-Free DNase Set (QIAGEN) according to the manufactureŕs instructions. Total RNA was DNase digested for 30 min at 37°C using 1 µl TURBO DNase (2 U/µl) (Invitrogen). A 500-ng aliquot of the isolated RNA was converted to cDNA with the iScript Advanced Kit (Bio-Rad) according to the manufacturer’s instructions. 1 µl of a 1:10 dilution in nuclease-free water of the cDNA samples was mixed with 5 µl of SsoAdvanced Univ SYBR Green Supermix (Bio-Rad) and 0.8 µl of 5 µM forward and reverse primers (Table S10). The total reaction volume was adjusted to 10 µl with nuclease-free water, dispensed in triplicates in a 96-well PCR plate (Bio-Rad) and subjected to qPCR in a Bio-Rad CFX real-time cycler. Evaluation of the obtained data was performed according to the ΔΔC_t_ method,^144^ using *recA* or *secA* genes as internal references.

#### Acid shock assay

Acid resistance was determined based on previously described protocols^15, 145^ with the following modifications: *E. coli* BW25113 cells were grown at 37 °C in LB pH 7.6 to OD_600_ = 0.5, adjusted to OD_600_ = 1, and then shifted for 15 min each first to LB pH 5.8 and then to LB pH 4.4. Then cells were shifted to LB pH = 3.0 for 1 h at 37°C. As a control, cells were cultivated at pH 7.6 throughout the experiment. After 1 h at pH 7.6, or pH 3, samples were serially diluted in 1x phosphate-buffered saline (PBS) and plated on LB agar plates to count the number of colonies. Percent survival was calculated as the ratio of colony forming units at pH 3 and pH 7.6.

#### Promoter activity assay

In vivo promoter activities of *gadBC*, *adiA* and *cadBA* were determined using luminescence-based reporter plasmids harboring fusions of the respective promoter regions to the *luxCDABE* genes from *Photorhabdus luminescens.* BW25113 wild-type cells or corresponding mutants from the Keio collection^83^ were transformed with plasmids with plasmid pBBR1-P*gadBC:lux,* pBBR1-P*adiA:lux* or pBBR1-P*cadBA:lux*. All strains were cultivated in LB medium supplemented with gentamycin overnight. The overnight cultures were inoculated to an OD_600_ of 0.05 in fresh LB medium (pH 7.6), aerobically cultivated until exponential phase (OD_600_ = 0.5) and shifted to LB pH 5.8. To assess promoter activity of *adiA* and *gadBC*, the cultures were shifted again to LB pH 4.4 after 15 min of growth in LB pH 5.8. In the next step, the cells were transferred to a 96-well plate and aerobically cultivated at 37°C in LB medium at different pH values supplemented with gentamycin. Growth and bioluminescence were measured every 10 min in the microtiter plates using a CLARIOstar Plus plate reader (BMG Labtech). Data are reported as relative light units (RLU) in counts per second of OD_600_.

#### Autoencoder-based identification of acid-specific genes and biological processes

The denoising autoencoders in this study were implemented using the Python package Keras.^146^ Details of the various hyperparameter and network architecture choices were taken from Kion-Crosby and Barquist 2023.^111^ In brief, an ensemble of 100 denoising autoencoders (DAEs) with two hidden layers of 2,000 and 1,000 nodes between the input layer and the bottleneck was used. The bottleneck layer of each network consists of 50 nodes as this was found to be within an optimal range for DAEs trained on the PRECISE 2.0 expression compendium^147^ by Kion-Crosby and Barquist 2023.^111^ All layers have sigmoid activation functions, the weights of each layer were randomly initialized based on the Glorot distribution, and the bias vectors with zeros. The weight matrices which make up the decoder of each network are tied such that they consist of the transpose of the corresponding weight matrices of each encoder.

All networks were trained in each ensemble using the Adam optimization algorithm. Data corruption during training was employed to improve generalizability such that 10% of the entries of each input data point were randomly set to zero during each training step. Additionally, early stopping was employed during training: the data were randomly portioned into an 80% training set and a 10% validation set, and training was stopped once the validation score began to worsen. A 10% test set was also portioned to determine the optimal training parameters. A local search over all training parameters including the learning rate and batch size was performed using the test score as a metric, and training was done with batch shuffling enabled.

For training of the autoencoder ensemble, the PRECISE 2.0 compendium^147^ was used as well as six additional data points of Bhatia *et al.* 2022^113^ representing *E. coli* K-12 MG1655 strains grown in M9 medium and treated with various antibiotics, in addition to the 9 data points from the current study. All data were converted to log transcripts per million, and features were normalized after the train/validation/test split such that the expression of each gene was scaled between 0 and 1.

Following the procedure described in Kion-Crosby and Barquist 2023,^111^ the encoder of each of the 100 trained networks was used to determine which nodes turn on for the specific stress conditions of interest (e.g. pH 4.4) and simultaneously turn off for alternative stress conditions (e.g. oxidative stress, heat stress, ethanol, etc.). This was done by passing each data point into each encoder while observing the activations of nodes at the bottleneck layer. After identifying which nodes are specific to the stress condition of interest, each corresponding decoder was utilized to generate gene expression predictions for all *E. coli* genes by activating each of these bottleneck nodes individually and propagating this signal through the decoder. After sorting all genes based on the average decoder output from all identified nodes, the top of this sorted vector was used to define a gene set associated with the condition of interest.

Additionally, GSEA was run on the outputs of each decoder for each node to determine which biological processes were associated with the condition of interest. Since these groups often consist of ∼100 nodes, and thousands of GO terms are evaluated, a conservative threshold for the adj. p-value of 0.0005 was set. The adj. p-value was found using the BH method, and each p-value was computed based on 1,000,000 permutations. Finally, for the selection of the high-confidence acid-specific gene candidates, first the log_2_FC between each acidic condition and every other stress condition was taken for all genes in the acid-specific gene sets. Genes corresponding to a log_2_FC of at least 0.5 for at least 95% of comparisons were then selected.

#### Propidium iodide viability staining

Cells were grown either at pH 7.6, 5.8, or 4.4 in the same manner as described for Ribo-Seq and RNA-Seq (Fig. 1A). After cultivation at the respective pH values, 1 ml of each culture was centrifuged (15,000 *g*, room temperature) and washed with PBS. Propidium iodide (Invitrogen) was added with a final concentration of 2 µg/ml, followed by 5 min incubation at room temperature in the dark to label dead cells. After another wash step using PBS, 2 μl of the culture was spotted on 1% (w/v) agarose pads, placed onto microscope slides and covered with a coverslip. Microscopic images were taken using a Leica DMi8 inverted microscope equipped with a Leica DFC365 FX camera. An excitation wavelength of 546 nm and a 605-nm emission filter with a 75-nm bandwidth was used to detect fluorescence with an exposure of 500 ms, gain 5, and 100% intensity in the Leica LAS X 3.7.4 software.

To quantify relative fluorescent intensities (RF) of single cells, phase contrast and fluorescent images were analyzed using the ImageJ^148^ plugin for MicrobeJ.^149^ Default settings of MicrobeJ were used for cell segmentation (Fit shape, rod-shaped bacteria) apart from the following settings: area: 0.1-max μm^2^; length: 1.2–5 μm; width: 0.1–1 μm; curvature 0.−0.15 and angularity 0.−0.25 for *E. coli* cells. In total, ≥1000 cells were quantified per strain and condition and the background of the agarose pad was subtracted from each cell per field of view. Cells with RF values ≥ 300 after subtraction of the background were considered as dead.

### QUANTIFICATION AND STATISTICAL ANALYSIS

One-way ANOVA followed by Bonferronís multiple comparisons tests were performed using GraphPad Prism version 8.4.3 for Windows. Differences were considered significant when p-values were less than 0.05. Pearson correlation coefficients were calculated using the *scipy.stats.pearsonr* function in Python 3.8.8. Mean and standard deviation are shown unless otherwise indicated in the figure legends. All experimental data are representative of at least three biological replicates.

### KEY RESOURCES TABLE

See separate word file.

#### Supplemental item titles

Figure S1. Related to Figure 1. Temporal dynamics of *adiA* transcription under acid stress (pH 4.4).

Figure S2. Related to Figure 1. Quantification of non-viable cells at varying pH conditions.

Figure S3. Related to Figures 1 and 2. Read mapping statistics for Ribo-Seq and RNA-Seq data.

Figure S4. Related to Figures 1 and 2. Read length distribution of ribosome-protected mRNA fragments (RPF) at different degrees of acidity.

Figure S5. Related to Figure 2. Reduction of ribosome occupancy in translation initiation regions at pH 5.8 and pH 4.4.

Figure S6. Related to Figure 2. Verification of differentially expressed genes under acid stress using RT-qPCR.

Figure S7. Related to Figure 4. Transcriptional and translational expression profiles of genes associated with enzyme-based H^+^-consuming acid resistance (AR) systems in *E. coli*.

Figure S8. Related to Figure 5. Subcellular location and homology of sORF15.

Table S1. Related to Figure 1. OD_600_ values determined at t_30_ (Fig. 1A) prior to sample collection for Ribo-Seq and RNA-Seq experiments.

Table S2. Related to Figure 1. pH values monitored during cultivation of *E. coli* for Ribo-Seq and RNA-Seq experiments (Fig. 1A).

Table S3. Related to Figure 2. Ribo-Seq and RNA-Seq exclusive genes.

Table S4. Related to Figure 2. Genes with differential translation efficiency under acid stress.

Table S5. Related to Figures 2, 4 and 5. Complete HRIBO results.

Table S6. Related to Figure 4. Candidate transcriptional regulators evaluated in Figure 4.

Table S7. Related to Figure 5. Acid-induced novel candidate sORFs.

Table S8. Related to Figure 5. Differential expression of sORFs under acid stress.

Table S9. Related to Figure 1 and 4. *Escherichia coli* strains used in this study.

Table S10. Related to Figures 4, 6 and S6. Oligonucleotides used in this study.

