## Supplementary information for "Ribosome profiling reveals the fine-tuned response of *Escherichia coli* to mild and severe acid stress"

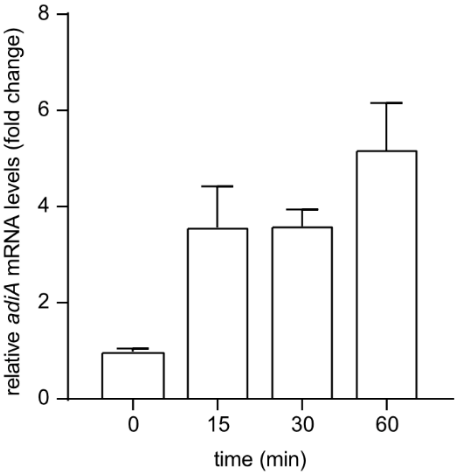


**Figure S1. Related to Figure 1.** Temporal dynamics of *adiA* transcription under acid stress (pH 4.4). Cells were cultivated as described in Figure 1. After the shift to pH 4.4, cells were collected after 0, 15, 30 and 60 min and total RNA was prepared. Relative levels of *adiA* mRNA were quantified by RT-qPCR. Fold-change values were determined relative to the 0-min time point and normalized using *recA* as a reference gene. Error bars indicate the standard deviation of three independent biological replicates (n=3).

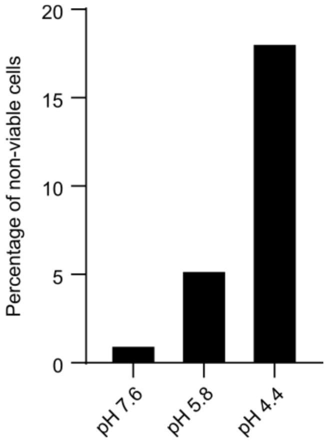


**Figure S2. Related to Figure 1.** Quantification of non-viable cells at varying pH conditions. *E. coli* MG1655 cells were cultured as outlined in Figure 1. Following sample collection, dead cells were distinguished by propidium iodide staining. Microscopy was performed using a Leica DMi8 inverted microscope equipped with a Leica DFC365 FX camera. A minimum of 1,000 cells were evaluated per condition, and relative fluorescence was measured using the MicrobeJ plugin for the ImageJ software. Cells exhibiting relative fluorescence values ≥300 after subtraction of the background were considered non-viable.

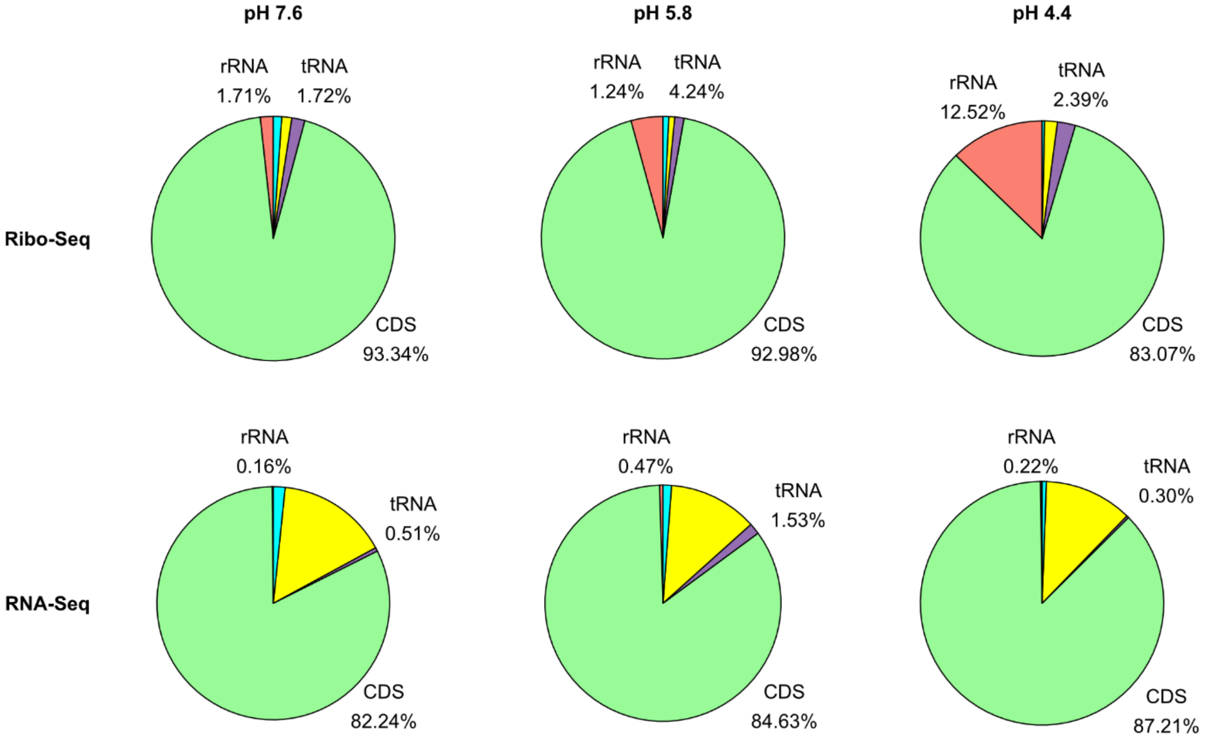


**Figure S3. Related to Figures 1 and 2.** Read mapping statistics for Ribo-Seq and RNA-Seq data. Pie charts illustrate the percentage of uniquely mapped reads either to CDS (green), rRNA (salmon), tRNA (purple), ncRNA (yellow), or pseudogenes (lightblue). The percentages provided indicate the average ratios relative to the total number of uniquely mapped reads per condition, which were calculated from biological triplicates. CDS, coding sequence; rRNA, ribosomal RNA; tRNA, transfer RNA; nc RNA, non-coding RNA.

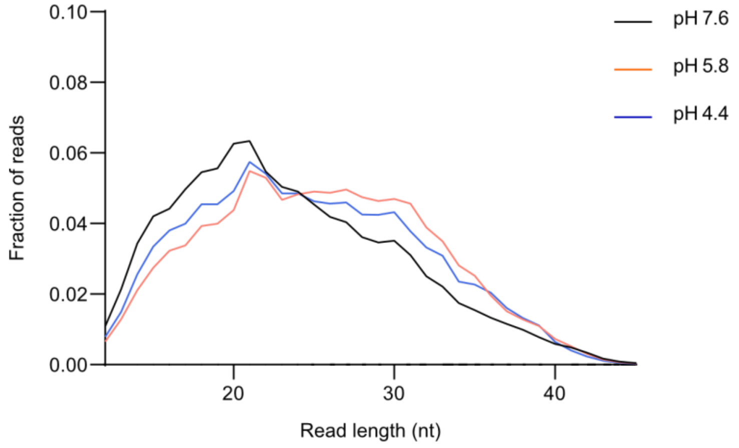


**Figure S4. Related to Figures 1 and 2.** Read length distribution of ribosome-protected mRNA fragments (RPF) at different degrees of acidity.

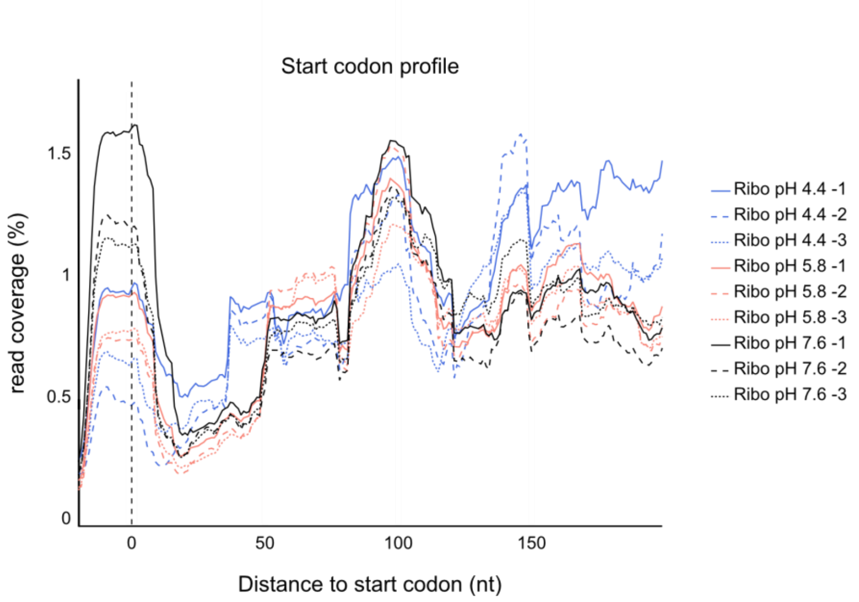


**Figure S5. Related to Figure 2.** Reduction of ribosome occupancy in translation initiation regions at pH 5.8 and pH 4.4. Alignments of reads from ribosome-protected fragments in the 5'UTR and the first 200 nucleotides of the coding sequence from all *E. coli* genes. The percentage of read coverage is shown relative to the total number of mapped reads per condition for each specific nucleotide position. The position of the first nucleotide of the start codon is indicated by a vertical dashed line.

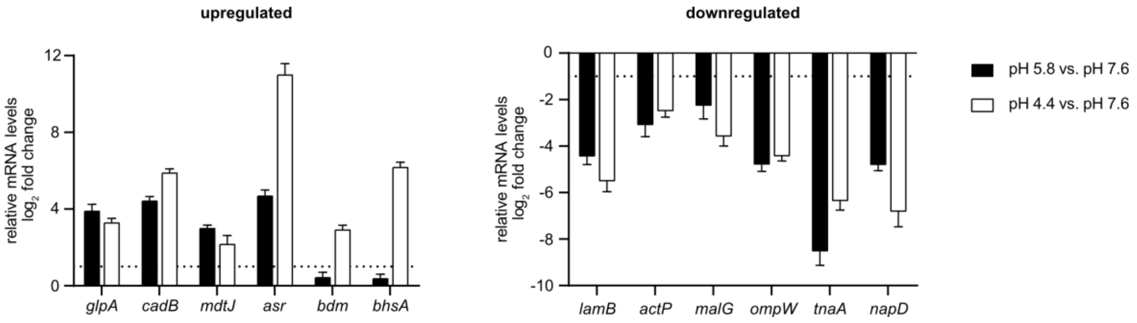


**Figure S6. Related to Figure 2.** Verification of differentially expressed genes under acid stress using RT-qPCR. Cells were cultivated as described in Figure 1. Relative mRNA levels were measured by RT-qPCR and fold change values were calculated relative to pH 7.6 and normalized using *recA* as a reference gene. Error bars indicate the standard deviation of three independent biological replicates (n=3).

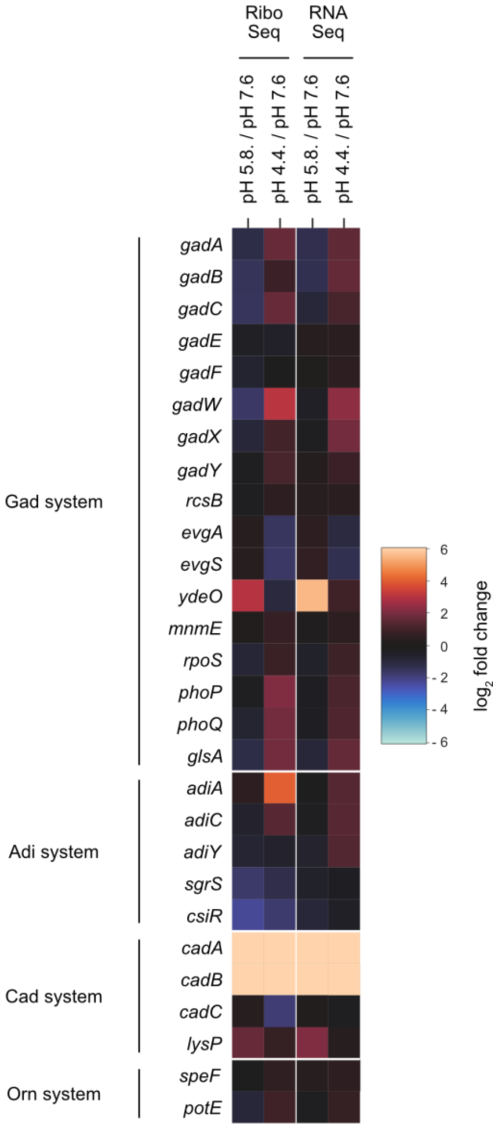


**Figure S7. Related to Figure 4:** Transcriptional and translational expression profiles of genes associated with enzyme-based $H^+$-consuming acid resistance (AR) systems in *E. coli*. Heatmap displaying RNA-Seq and Ribo-Seq $log_2$ fold change values of genes encoding amino acid decarboxylases, antiporters, or regulatory elements associated with either the Gad, Adi, Cad, or Orn system. $Log_2$ fold changes of normalized expression values at pH 4.4 and 5.8 were calculated relative to the normalized expression values at pH 7.6.

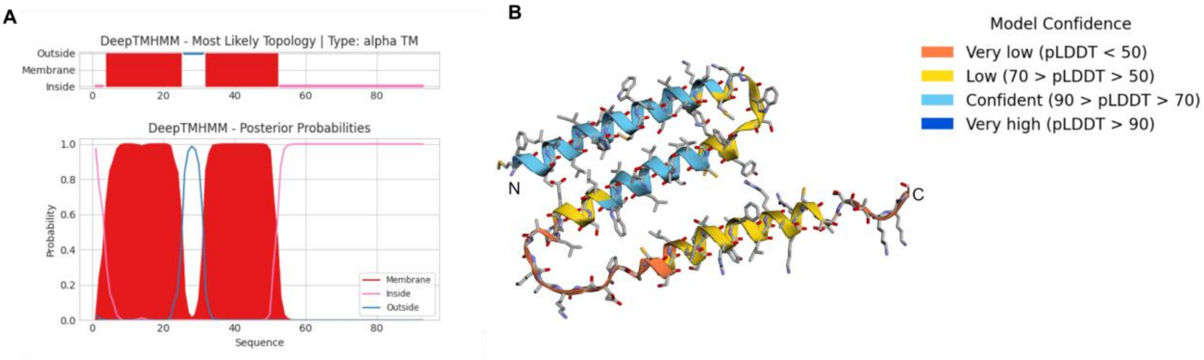


C

| Species | Min E-value | Max %identity | Annotation |
|---|---|---|---|
| *Vibrio parahaemolyticus* | 4.48E-39 | 100 | hypothetical protein |
| *Shigella flexneri* | 2.46E-27 | 100 | hypothetical protein |
| *Klebsiella pneumoniae* | 3.42E-44 | 98.67 | hypothetical protein |
| *Salmonella enterica* | 2.14E-41 | 98.61 | hypothetical protein |
| *Shigella boydii* | 7.4E-30 | 98.21 | hypothetical protein |
| *Shigella sonnei* | 1.98E-56 | 97.85 | hypothetical protein |
| *Enterococcus faecium* | 1.06E-37 | 97.01 | hypothetical protein |
| *Escherichia albertii* | 2.57E-43 | 88.1 | hypothetical protein |

**Figure S8. Related to Figure 5:** Subcellular location and homology of sORF15. **(A)** Transmembrane topology of sORF15 predicted by DeepTMHMM. (B) Predicted protein structure of sORF15 using ColabFold. N- and C-termini are indicated **(C)** Homologs of sORF15 identified with blastp. The sORF15 amino acid sequence was used as the query sequence and homologs with an E-value < 0.05 and Max % identity of > 80% are listed.

**Table S1. Related to Figure 1.** $OD_{600}$ values determined at $t_{30}$ (Fig. 1A) prior to sample collection for Ribo-Seq and RNA-Seq experiments.

| | Replicate I | | | Replicate II | | | Replicate III | | |
|---|---|---|---|---|---|---|---|---|---|
| | pH 7.6 | pH 5.8 | pH 4.4 | pH 7.6 | pH 5.8 | pH 4.4 | pH 7.6 | pH 5.8 | pH 4.4 |
| $OD_{600}$ ($t_{30}$) | 1.05 | 0.93 | 0.69 | 1.18 | 1.08 | 0.66 | 1.13 | 1.04 | 0.72 |

**Table S2. Related to Figure 1.** pH values monitored during cultivation of *E. coli* for Ribo-Seq and RNA-Seq experiments (Fig. 1A). pH-shifts were initiated by direct addition of 5 M HCl to the cultures and are indicated by (*****).

| | Replicate I | | | Replicate II | | | Replicate III | | |
|---|---|---|---|---|---|---|---|---|---|
| | pH 7.6 | pH 5.8 | pH 4.4 | pH 7.6 | pH 5.8 | pH 4.4 | pH 7.6 | pH 5.8 | pH 4.4 |
| pH 0 min ($t_0$) | 7.24 | 5.90***** | 5.84***** | 7.29 | 5.84***** | 5.82***** | 7.28 | 5.81***** | 5.87***** |
| pH 15 min ($t_{15}$) | 7.17 | 5.95 | 4.41***** | 7.21 | 5.87 | 4.37***** | 7.24 | 5.83 | 4.33***** |
| pH 30 min ($t_{30}$) | 7.15 | 6.01 | 4.43 | 7.17 | 5.93 | 4.39 | 7.16 | 5.88 | 4.39 |

**Table S6. Related to Figure 4.** Candidate transcriptional regulators evaluated in Figure 4. Transcription factors were chosen based on Ribo-Seq $log_2$ fold change values at either pH 4.4 or pH 5.8 compared to pH 7.6.

| Regulator | $log_2$ fold change |
|---|---|
| YdeO | 2.96 (pH 5.8) |
| MhpR | 4.72 (pH 4.4) |
| IscR | 4.52 (pH 4.4) |
| MarR | 3.10 (pH 4.4) |
| GadW | 3.03 (pH 4.4) |
| YdcI | 2.92 (pH 4.4) |
