## Supplementary material for "Ribosome profiling reveals the fine-tuned response of *Escherichia coli* to mild and severe acid stress": Key resources table

| REAGENT or RESOURCE | SOURCE | IDENTIFIER |
|---|---|---|
| **Bacterial and Virus Strains** | | |
| | See Table S9 | |
| **Chemicals, Peptides, and Recombinant Proteins** | | |
| Micrococcal Nuclease (MNase) | New England BioLabs | Cat# M0247S |
| SYBR Gold | Invitrogen | Cat# S11494 |
| T4 Polynucleotide Kinase | New England BioLabs | Cat# M0201S |
| Adenosine 5´Trophosphate (ATP) | New England BioLabs | Cat# P0756S |
| Propidium iodide | Invitrogen | Cat# P3566 |
| NEBuilder HiFi DNA Assembly | New England BioLabs | Cat# E2621L |
| SsoAdvanced Univ SYBR Green Supermix | Bio-Rad | Cat# 1725271 |
| TURBO DNase | Invitrogen | Cat# AM2238 |
| **Critical Commercial Assays** | | |
| miRNeasy Mini Kit | QIAGEN | Cat# 217004 |
| RNase-Free DNase Set | QIAGEN | Cat# 79254 |
| RNA MinElute Cleanup Kit | QIAGEN | Cat# 74204 |
| QIAquick PCR Purification Kit | QIAGEN | Cat# 28104 |
| NEBNext Small RNA Library Prep Set for Illumina | New England BioLabs | Cat# E7580S |
| NEBNext Ultra II Directional RNA Library Prep Kit for Illumina | New England BioLabs | Cat# E7760L |
| NEBNext rRNA Depletion Kit (Bacteria) | New England BioLabs | Cat# E7850L |
| RNA 6000 Nano Kit | Agilent | Cat# 5067-1511 |
| High Sensitivity DNA Kit | Agilent | Cat# 5067-4626 |
| Qubit RNA HS Assay Kit | Invitrogen | Cat# Q32855 |
| Quick-RNA Miniprep Kit | Zymo Research | Cat# R1055 |
| iScript Advanced cDNA Synthesis Kit | Bio-rad | Cat# 1725038 |
| **Deposited Data** | | |
| RIBO-Seq data | Gene Expression Omnibus (GEO) | GSE219022 |
| RNA-Seq data | Gene Expression Omnibus (GEO) | GSE219022 |
| **Oligonucleotides** | | |
| | See Table S10 | |
| **Recombinant DNA** | | |
| pBBR1-MCS5-P*gadBC:lux* | This study | N/A |
| pBBR1-MCS5-P*adiA:lux* | Brameyer et al. 2022 | N/A |
| pBBR1-MCS5-P*cadBA:lux* | Brameyer et al. 2020 | N/A |
| **Software and Algorithms** | | |
| HRIBO 1.6.0 | Gelhausen et al. 2021 | https://github.com/RickGelhausen/HRIBO |
| Python 3.8.8 | Python Software Foundation | https://www.python.org/ |
| jbrowse2 2.2.2 | Diesh et al. 2022 | https://jbrowse.org/jb2/ |
| plotly 5.11.0 | Plotly Technologies Inc | https://plot.ly |
| Rscript 4.1.3 | R Core Team 2021 | https://www.R-project.org |
| DESeq2 1.38.0 | Love et al. 2014 | https://github.com/mikelove/DESeq2 |
| clusterprofiler 4.2.0 | Wu et al. 2021 | https://github.com/YuLab-SMU/clusterProfiler |
| GraphPad Prism version 8.4.3 for Windows | GraphPad Software, San Diego, California, USA | https://www.graphpad.com/ |
| CLC Main Workbench 20.0.4 | QIAGEN | https://digitalinsights.qiagen.com/ |
| Microbe J 5.13l | Ducret, Quardokus and Brun, 2016 | https://www.microbej.com/ |
| LAS X 3.7.4 | Leica | https://www.leica-microsystems.com/ |
| PSORTb 3.0.3 | Yu et al. 2010 | https://www.psort.org/psortb/ |
| DeepTMHMM 1.0.24 | Hallgren et al. 2022 | https://dtu.biolib.com/DeepTMHMM/ |

| | | |
|---|---|---|
| Keras | Cholet 2015 | https://keras.io. |
